# Spatial and Temporal Cortical Variability Track with Age and Affective Experience During Emotion Regulation in Youth

**DOI:** 10.1101/291245

**Authors:** João F. Guassi Moreira, Katie A. McLaughlin, Jennifer A. Silvers

## Abstract

Variability is a fundamental feature of human brain activity that is particularly pronounced during development. However, developmental neuroimaging research has only recently begun to move beyond characterizing brain function exclusively in terms of magnitude of neural activation to incorporate estimates of variability. No prior neuroimaging study has done so in the domain of emotion regulation. We investigated how age and affective experiences relate to spatial and temporal variability in neural activity during emotion regulation. In the current study, 70 typically developing youth aged 8-17 years completed a cognitive reappraisal task of emotion regulation while undergoing functional magnetic resonance imaging. Estimates of spatial and temporal variability during regulation were calculated across a network of brain regions, defined *a priori*, and were then related to age and affective experiences. Results showed that increasing age was associated with reduced spatial and temporal variability in a set of frontoparietal regions (e.g., dorsomedial prefrontal cortex, superior parietal lobule) known to be involved in effortful emotion regulation. In addition, youth who reported less negative affect during regulation had less spatial variability in the ventrolateral prefrontal cortex, which has previously been linked to cognitive reappraisal. We interpret age-related reductions in spatial and temporal variability as implying neural specialization. These results suggest that the development of emotion regulation is undergirded by a process of neural specialization and open up a host of possibilities for incorporating neural variability into the study of emotion regulation development.

---

Advances in *in vivo* functional neuroimaging have granted psychologists novel insights into how the human brain develops and functions. While most research in developmental neuroscience has focused on comparing the *magnitude* of neural responses at different ages, accumulating evidence suggests that it is also important to assess neural *variability* – that is, how the neural signal varies across time and brain region within and between individuals (e.g., Durston et al., 2006; Heller & Casey, 2016; Nomi, Bolt, Ezie, Uddin, & Heller, 2017). Variability is another dimension by which to categorize brain function and age-related differences in neural variability likely reflect important developmental processes, including the degree of specialization and experience-based plasticity (e.g., pruning) in neural circuits across age (Casey, 2015; Durston et al., 2006). Emotion regulation presents itself as a particularly important skill to be assessed in a developmental neural variability framework because it exhibits protracted maturation and is critical for wellbeing (Cole & Deater-Deckard, 2009; Gross, 2015; McLaughlin, Garrard, & Somerville, 2015). However, virtually all developmental neuroimaging studies of emotion regulation to date have concentrated on age-related differences in the magnitude of brain activity *across* individuals and have ignored how variability *within* individuals during emotion regulation differs across development. The present study sought to address this knowledge gap by investigating how within-subject variability in neural activity during emotion regulation—both spatial and temporal—is associated with age and affective experience in youth.

## Conceptualizing Emotion Regulation and Clarifying Theoretical Stances

Emotion regulation, frequently defined as modulation of affective processes in accordance with explicit or implicit goals (Etkin, Büchel, & Gross, 2015), requires years of development before maturity is reached (Guassi Moreira & Silvers, 2018). Over the past three decades, an explosion of theoretical and empirical advances has helped to establish various frameworks for studying emotion regulation (Camras, 2011; Cole, Martin, & Dennis, 2004; Gross & Feldman Barrett, 2011). Across diverse theoretical orientations, it is largely accepted that emotion regulation is dynamic, occurs over multiple timescales, and draws upon a variety of supporting psychological processes (Cole et al., 2004; Etkin et al., 2015; Gross & Feldman Barrett, 2011; Ochsner, Silvers, & Buhle, 2012). The dynamic nature of emotion regulation requires that developmental research on this topic consider age-related changes to the contexts in which regulation occurs and the bottom-up affective experiences that trigger regulation. Put another way, the very emotion that is being regulated likely changes with age at the same time that regulatory abilities change. Moreover, given seminal research demonstrating that emotional experiences change not only in their means but also in their variability across age (Larson, Csikszentmihalyi, & Graef, 1980), this work signals the importance for focusing on variability in developmental emotion regulation research.

While the experimental paradigm used in this manuscript conforms most directly to appraisal theory (Gross, 2015), considering the role of neural variability in neurodevelopment stands to inform a number of theoretical approaches to the study of emotion regulation. The purpose of this report is not necessarily to support or refute one particular theoretical ideology; instead we outline several examples throughout for how many different theoretical views might stand to benefit by considering variability.

## Prior Investigations of Emotion Regulation Neurodevelopment Omit Variability

Inspired by behavioral work demonstrating that emotion regulation abilities improve significantly during childhood and adolescence (Kim & Richardson, 2010; Silvers et al., 2012; Somerville, Jones, & Casey, 2010; Thompson & Goodman, 2010), a growing number of neuroimaging studies have begun to explore the neural bases of emotion regulation in youth. One popular means for doing so has been to employ functional magnetic resonance imaging (fMRI) in conjunction with tasks examining cognitive reappraisal—a widely studied and adaptive emotion regulation strategy that involves thinking about a stimulus differently in order to modulate its emotional import (Denny & Ochsner, 2014; Giuliani & Pfeifer, 2015; Gross, 2015; Ochsner, Silvers, & Buhle, 2012). In healthy adults, reappraisal recruits frontoparietal regions commonly implicated in cognitive control, including the ventrolateral, dorsolateral and dorsomedial prefrontal cortex (vlPFC, dlPFC, and dmPFC, respectively) as well as superior parietal lobule (SPL; Buhle et al., 2014; Ochsner et al., 2004, 2012; Ochsner & Gross, 2005). Across development, increasing age is associated with increased recruitment of these frontoparietal regions along with reduced negative affect, suggesting that emotion regulation success improves as cognitive control abilities become more fine-tuned (McRae et al., 2012; Silvers et al., 2012, 2016; Silvers, Shu, Hubbard, Weber, & Ochsner, 2015). These age-related neural and behavioral changes are observed when individuals are instructed to reappraise but not when instructed to react naturally to emotional stimuli (McRae et al., 2012; Silvers et al., 2012, 2016; Silvers, Shu, Hubbard, Weber, & Ochsner, 2015), suggesting that changes in emotional responding are driven more strongly by changing regulatory abilities than by changing baseline emotional experience. While informative, these initial developmental neuroimaging studies of reappraisal have relied heavily on univariate analyses that characterize age-related changes in terms of peak or mean BOLD signal. As such, this existing work has helped advance the field, but has also overlooked the role that neural variability may play in emotion regulation.

## Variability is a Key Feature of Neural Function and a Catalyst for Development

Variability is a fundamental feature of brain activity that is distinct from “randomness” or “noise” (Pinneo, 1966). Neural activity is organized according to structured spatial and temporal profiles (Christophel, Iamshchinina, Yan, Allefeld, & Haynes, 2018; Huth, Heer, Griffiths, Theunissen, & Gallant, 2016; Huth, Nishimoto, Vu, & Gallant, 2012; Luciana, Wahlstrom, Porter, & Collins, 2012). While not always thought of in terms of variability, the fact that spatial and temporal activation patterns within-individuals vary between different psychological processes suggests that variability is a defining feature of brain function (Etzel, Zacks, & Braver, 2013; Patel, Kaplan, & Snyder, 2014). Neuronal computations vary for different psychological processes, leading neuroimaging data associated with different processes to vary as well (Kriegeskorte, Cusack, & Bandettini, 2010). The mere fact that seeing a face evokes a different pattern of activity than seeing a house, for example, illustrates that configurations of brain activity are spatially variable (Haxby et al., 2001). Examining variability is therefore one meaningful way to characterize the neural substrates of psychological processes. We can think of three diverse reasons, encompassing both psychological and neuroscientific perspectives, for why this notion matters. First, variability is another fundamental dimension—in addition to magnitude—along which psychological processes are embedded in the neural code. If scientists want to understand the totality of how the brain gives rise to psychology, it would be advantageous to examine this dimension as the neural underpinnings of psychological phenomena may also be encoded in terms of variability (in addition to magnitude). In doing so, researchers can quantify and parameterize developmental change in a previously overlooked arena to glean novel insights. Second, prior neuroscience research has revealed that subtle fluctuations in neural activity are often more informative of psychological processes than peak activations (e.g., Clare Kelly, Uddin, Biswal, Castellanos, & Milham, 2008; Nomi, Bolt, Ezie, Uddin, & Heller, 2017; Raichle et al., 2001). While the same is likely true for emotion regulation, this possibility remains untested. Last, developmental psychology research highlights that mean affective states change less across age when compared to affective variability (Larson, Csikszentmihalyi, & Graef, 1980; Larson & Lampman-Petraitis, 1989; Silvers et al., 2012). However, neuroscientific research has yet to address this point.

Though within-individual variability pertaining to the neurodevelopment of emotion regulation has not been examined, prior research in psychology and developmental neuroscience can help scaffold the present investigation. Psychologically, children try a variety of strategies during development (i.e., exhibit high variability) before becoming expert in a narrower set of strategies (Roalf et al., 2014; Siegler, 1994, 2007). Brain development appears to follow a similar pattern. Functionally, brain responses may exhibit “focalization” across age – a shift from a more variable spatial signature of activity to one that is more concentrated, and potentially, specialized (Dehaene-Lambertz, Monzalvo, & Dehaene, 2018; Durston et al., 2006; Richardson, Lisandrelli, Riobueno-Naylor, & Saxe, 2018). Importantly, however, evidence for the shift from diffuse to focused activity has come from studies examining average levels of brain activity across individuals of different ages rather than examining within-subject variability. These patterns of functional maturation may be driven, in part, by pruning of initially over-produced synaptic connections to produce increasingly specialized neural networks that retain only their most essential connections (DuPre & Spreng, 2017; Durston et al., 2006; Foulkes & Blakemore, 2018; Sowell, Thompson, Tessner, & Toga, 2001; Vij, Nomi, Dajani, & Uddin, 2018). These findings suggest that neurodevelopment is characterized by a transition from diffuse and spatially variable patterns of activity towards optimized and constrained functional networks.

Investigating neural variability promises to enhance understanding of neurodevelopment and generate novel hypotheses for future research (Poldrack, 2015). For instance, studying within-person variability can identify another dimension by which the activity of brain regions and networks differs or co-varies. This could help to categorize emotion regulation strategies according to underlying mental structure and to examine how psychological subprocesses associated with different forms of emotion regulation develop across age (Braunstein et al., 2017; Eisenberg et al., 2018). Relatedly, variability may serve as another metric for assessing maturation of emotion regulation. As an example, the dynamic systems view of development posits that variability within individuals is a hallmark of developmental processes because systems are always ‘in flux’, and that different developmental stages are marked by different patterns of variability (see Box 1 of Smith & Thelen, 2003). This variability can manifest at the neural level, as we examine in this report, as well as the behavioral level (e.g., youth trying variations in emotion regulation tactics until settling upon a preferred strategy). Under this view, formally quantifying variability might allow for more precise mapping of individual brain development growth curves as it relates to emotion regulation skills across age in addition to charting what is the most adaptive and ideal amount of variability for a given system across development.

## Incorporating Variability into the Study of Emotion Regulation Neurodevelopment

We used the work summarized above to guide our query into how neural variability supports the neurodevelopment of emotion regulation. In this study, we specifically focus on spatial and temporal variability.

*Spatial Variability.* Spatial variability, or differences in how activation is distributed across a brain region, is an important organizational feature of neural activity. Psychological processes, including emotional experiences, are encoded in richly detailed spatial topographies that blanket the cortical landscape (Chang, Gianaros, Manuck, & Krishnan, 2015; Huth et al., 2016, 2012; Kriegeskorte et al., 2008). Prior work in developmental neuroscience suggests that such topographies are diffuse earlier in life and become focalized with age and experience (Dehaene-Lambertz et al., 2018; Durston et al., 2006; Park et al., 2004; Polk et al., 2002). Examining spatial variability during a specific psychological process may be one way to infer how specialized a given brain region is for that process for a given individual. For example, if an individual shows a low degree of spatial variability within ventrolateral prefrontal cortex (vlPFC) during emotion regulation, this would suggest that activity is concentrated to a specific subset of the neurons in that region. In contrast, a high degree of spatial variability in another individual would suggest that the computations occurring in vlPFC that support emotion regulation are being carried out across a larger population of neurons for that person. Examining spatial variability promises to uncover insights about the functional architecture of neurodevelopment that has relevance not only for emotion regulation but for many other psychological processes. In the current study, we implemented a novel method of estimating spatial variability by repurposing an analytic technique from economics—Gini coefficients—to serve as a metric of spatial variability in a given brain region. As will be described later, Gini coefficients are an especially useful tool because they can yield a numerical index of spatial variability, helping to quantify complex theoretical concepts such as focalization.

This approach can be particularly informative for affective neuroscience by way of its ability to highlight whether certain brain regions co-vary or differ in their topographic organization by telling us how patterns of activity are arranged across the surface of cortex (i.e., concentrated or diffusely). For instance, vlPFC, dlPFC, and dmPFC may all show similar magnitudes of activity during reappraisal, but vlPFC and dlPFC may have more similar topographies (i.e., both diffuse or both focal), indicating that a different psychological sub-process is being implemented by those brain regions compared to the computations in dmPFC (e.g., working memory vs mentalizing). One potential theoretical application of this work would be to help test dominant theories that posit multiple executive functions support complex, cognitively demanding emotion regulation strategies (Ochsner et al., 2012).

*Temporal Variability.* While spatial variability can describe how brain function is organized topographically, temporal variability can lend insight into how brain activity changes over time to meet the dynamic demands of one’s environment. This is especially relevant for psychological phenomena that change over time, such as emotion regulation (Aldao, Sheppes, & Gross, 2015; Heller & Casey, 2016). Studying temporal variability in brain activation across multiple instances of emotion regulation can lend insight into how neural computations adjust according to varying regulatory demands. Situations that require emotion regulation vary markedly – emotion regulation may be required to maintain calm in a traffic jam or to respond to the loss of a loved one. Learning to consistently mount an effective regulatory response to variable affective events is a significant developmental hurdle. Given that youth tend to experience more intense and labile affect, and yet have fewer cognitive resources to draw from, we might expect them to display less consistent (i.e., more variable) neural recruitment during self-regulation (Larson, Csikszentmihalyi, & Graef, 1980; Mischel & Mischel, 1983; Silvers et al., 2012; Somerville et al., 2010). As individuals mature and experience with emotion regulation grows, however, neural computations underlying emotion regulation are likely to become more consistent (i.e., less variable; Koolschijn, Schel, de Rooij, Rombouts, & Crone, 2011; Ordaz, Foran, Velanova, & Luna, 2013).

## Current Study

In the current study, we used functional magnetic resonance imaging (fMRI) to examine spatial and temporal variability of frontoparietal brain responses during cognitive reappraisal—one particular emotion regulation strategy—in a sample of typically developing youth. We further sought to characterize how spatial and temporal variability related to age and affective experiences during emotion regulation via reappraisal in this sample. By doing so, we were able to observe how two forms of within-individual neural variability related to age and experiences of affect during emotion regulation (Foulkes & Blakemore, 2018). Given the paucity of research on this matter, this research was exploratory and guided by open questions regarding neural variability and emotion regulation rather than formal hypotheses. The methods and framework for studying variability described here are not necessarily beholden to one particular theoretical orientation, but rather showcase that variability is a meaningful feature of neurodevelopment with clear implications for a variety of perspectives on emotion regulation.

## Methods

*Participants.* Participants included 70 youth (34 female) ranging in age from 8.08 to 17.00 years (M*age* = 12.70). These participants were drawn from a larger sample taking part in a longitudinal study aimed at understanding the effects of childhood maltreatment on affective neurodevelopment. Only youth from the non-maltreated community control group were selected for the current analyses. Exclusion criteria for this group included exposure to childhood maltreatment or violence, presence of a developmental disorder (e.g., autism), psychotropic medication use, and IQ < 75. To qualify for inclusion in the current analysis, participants had to have: (i) accompanying behavioral data from the in-scan emotion regulation task (described below); (ii) low levels of motion during the scan (see below); and (iii) anatomical images that were free of abnormalities or artifacts. Participants and their families were recruited from a large, metropolitan area in the Pacific Northwest. Parents provided written consent and children provided written assent in accordance with the University of Washington Institutional Review Board. Participants were compensated $75 for completing the brain scan.

*fMRI Paradigm.* During the fMRI scan, participants completed a computerized version of a cognitive reappraisal task adapted from prior developmental studies of emotion regulation (McLaughlin, Peverill, Gold, Alves, & Sheridan, 2015; Silvers et al., 2016). Though youth have many emotion regulation strategies at their disposal (Braunstein, Gross, & Ochsner, 2017; Guassi Moreira & Silvers, 2018), a cognitive reappraisal task was chosen for a number of reasons. It is one of the most widely studied emotion regulation strategies in adult, pediatric, and clinical samples and thus had a robust literature to reference. It is frequently linked to a variety of adjustment outcomes (Denny & Ochsner, 2014; Giuliani & Pfeifer, 2015; Haines et al., 2016; Panno, Lauriola, & Figner, 2013). Last, it is more feasible to implement in a scanner environment compared to other emotion regulation strategies (e.g., situation selection or social support seeking). With this said, we note that the methods of estimating variability outlined below are not limited for use solely in the context of cognitive reappraisal and can be employed with other emotion regulation paradigms as well as a broader class of psychological processes altogether.

During the task, participants viewed a series of developmentally appropriate aversive and neutral images modeled after the International Affective Picture System (IAPS) stimulus set (Lang, Bradley, & Cuthbert, 2008). Great care was taken to create and validate a stimulus set that would be appropriate to use in children and adolescents. Aversive images all depicted conflict between individuals or personal struggles (e.g., an individual sitting alone in sadness). Moreover, such images were especially relevant for youth because they portrayed youth in aversive scenarios common to their lives (e.g., depiction of bullying/fighting behaviors, adults fighting in front of children). Images were purchased from a stock photography website (https://www.shutterstock.com), including 225 negative and 150 neutral images. In a pilot study, 120 children aged 6-16 years (50% female) provided ratings of valence, arousal, and dominance for a randomized selection of 80 images using the standardized assessment mannequins used to norm the IAPS stimuli. The stimuli and normative ratings are available on the lab website of the principal investigator of the original study (www.stressdevelopmentlab.org). Images were selected from the larger stimulus set based on the valence ratings from the pilot study; the images with the most negative valence ratings were selected for the negative condition, and those closest to the mid-point of the valence scale were selected for the neutral condition. This custom image validation procedure is a notable strength, as most developmental imaging investigations of emotion regulation typically rely on stimulus sets that have been developed and validated for use in adults.

During the task, participants were instructed to either passively observe or reappraise the images via psychological distancing. Afterwards, participants provided their ratings of negative affect along a four-point Likert scale (see Figure 1). Though there are limitations of self-report (reviewed in the discussion), it also has several strengths: (i) it is tightly associated with various physiological measures (Levesque et al., 2004; Ochsner & Gross, 2008; Ray, McRae, Ochsner, & Gross, 2010), (ii) it provides relatively direct access to one’s emotional experiences in ways that observational measures (including physiology) cannot (Larsen & Prizmic-Larsen, 2006), (iii) it is easily compatible with the scanning environment and offers a low bar in terms of task demands, and (iv) it is unambiguous relative to certain physiological measures such as heart rate variability (HRV) or respiratory sinus arrhythmia (RSA) (Heathers & Goodwin, 2017) where two or more psychological and affective signals can yield the same pattern of results.

**Figure 1.**
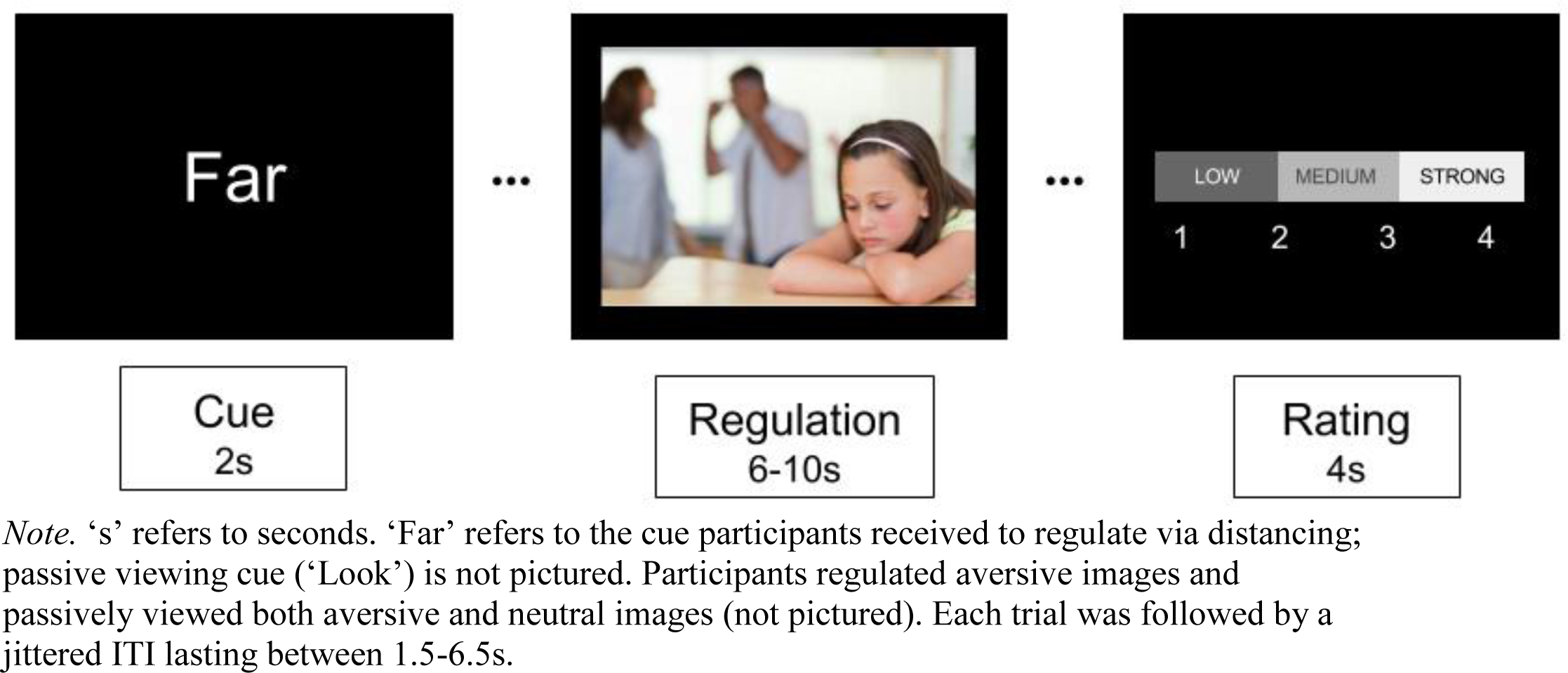
Overview of Emotion Regulation Paradigm

Participants reappraised negative stimuli on 20 trials, passively observed negative stimuli on 20 trials, and passively observed neutral stimuli on 20 trials. Prior to each image, an instructional cue was displayed that informed participants whether they were to passively observe (‘Look’) or regulate (‘Far’) the subsequent image. For ‘Look’ trials, participants were told to passively observe the stimulus and react to it as they normally would. During ‘Far’ trials, participants were trained to reappraise the image in a way that made it psychologically distant, such as pretending they were physically standing far from the image in each scene or that they were behind a movie camera, recording the events show in the picture. Since our interests lay in characterizing spatial and temporal variability during active emotion regulation, we focused on ‘Far’ trials for the current report.

Prior to scanning, participants completed a training session where they received information about the meaning of the instructional cues and how they were to think about stimuli presented after each type of cue. Experimenters then shared several examples of each condition to participants before asking them to complete a series of 5 practice trials for each condition.

During the in-scan task instructional cues were presented for 2 seconds and stimuli were jittered such that they were displayed for 6s – 10s. The Likert scale was presented thereafter for 4 seconds followed by a 1.5s – 6.5s ITI. Participants completed four runs, each lasting approximately 220s (runs ranged between 110 and 115 volumes in length). The stimuli used for ‘Look’ and ‘Far’ trials did not differ in their normative ratings of valence and arousal. The task was programmed using E-Prime (Psychology Software Tools, Inc., http://www.pstnet.com).

### fMRI Data Acquisition and Preprocessing

*fMRI Data Acquisition.* Prior to image acquisition, participants younger than 12 years old or who exhibited any signs of nervousness about scanning were taken to a mock MRI scanner to become familiarized with the scanning environment and trained on how to minimize head motion. These participants watched a film on a back-projector with a head-mounted motion tracker. The film paused if a head movement exceeding 2mm occurred, helping participants quickly learn to keep still while in the mock scanner bore. In addition to this measure, participants were packed into the head coil with an inflated, head-stabilizing pillow to restrict movement.

Image acquisition was conducted on a 3T Phillips Achieva scanner at the University of Washington (Seattle) Integrated Brain Imaging Center. A 32-channel head coil was implemented, along with the use of a parallel image acquisition system. T1-weighted, magnetization-prepared rapid acquisition gradient echo (MPRAGE) volumes were acquired (TR = 2530ms, TE = 1640-7040μs, 7º flip angle, 256 mm^2^ FOV, 176 slices, 1 mm^3^ isotropic voxel size). Blood oxygenation level dependent (BOLD) signal during functional runs was acquired using a gradient-echo T2*-weighted echoplanar imaging (EPI) sequence. Thirty-two 3-mm thick slices were acquired parallel to the AC-PC line (TR = 2000ms, TE = 30ms, 90º flip angle, 256 × 256 FOV, 64 × 64 matrix size).

*fMRI Data Pre-Processing.* Prior to pre-processing, data were visually inspected for artifacts and anatomical abnormalities. fMRI data were pre-processed and analyzed using the fMRI Expert Analysis Tool (FEAT, version 6.00) of the FMRIB Software Library package (FSL, version 5.0.9; fsl.fmrib.ox.ac.uk). Pre-processing consisted of using the brain extraction tool (BET) to remove non-brain tissue from functional and structural runs, spatial realignment of functional volumes to correct for head motion using MCFLIRT, and hi-pass filtering the data (100s cutoff). The extent of participant head motion was further estimated by running FSL Motion Outliers to record volumes that exceeded a 0.9 mm threshold of framewise displacement (FD) (Siegel et al., 2014). Runs were discarded if participants exceeded this threshold for more than 25 volumes (~20% of a single run). Six participants had at least one run discarded due to excessive head motion (Mean number of discarded runs for eligible participants = 2 runs). The average number of volumes exceeding our FD threshold per run, per participant was 2.333 (*SD* = 4.04, range = 0-16.5); prior to discarding runs it was 3.014 (*SD* = 6.09, range = 0 - 30.75). Since a goal of the study was to examine how spatial variability—both between subjects and across time within subjects—related to age and experiences of negative affect, we elected not to spatially smooth our data. Data were pre-whitened to account for autocorrelated residuals across time. In addition to spatial and temporal variability analyses described below, we also ran traditional univariate analyses. A description of preprocessing for said analyses and their results can be found in the Supplement.

*ROI Definition.* We identified seven brain regions implicated in emotion regulation from a prior meta-analysis (Buhle, Silvers, et al., 2014). For the rest of this report, we refer to these seven regions as “ROIs”. Each ROI from the original meta-analysis contained one global maximum—the peak voxel from the cluster—and at least one other local maximum (voxels that constitute regional peaks within the cluster). Global and local maxima from these ROIs are described as “spheres”.

We defined ROIs based on a multi-kernel density meta-analysis of cognitive reappraisal fMRI studies (Buhle, Silvers, et al., 2014) and created spheres around the maxima in each ROI (global and local). ROIs were identified based on clusters reported in Table 2 of Buhle, Silvers, et al., (2014), resulting in 7 ROIs: right dorsolateral prefrontal cortex (R dlPFC), left dorsolateral prefrontal cortex (L dlPFC), right ventrolateral prefrontal cortex (R vlPFC), right dorsomedial prefrontal cortex (R dmPFC), left middle temporal gyrus (L MTG), left superior parietal lobule (L SPL), and right superior parietal lobule (R SPL). We then drew spherical masks (4mm radius) around each maxima (global and local). On the rare occasions that spheres extended beyond the boundaries of the brain, spheres were moved inward. Our choice to use 4mm radius spheres was motivated by the fact that anything smaller would have rendered an insufficient number of voxels with which to calculate Gini coefficients, and that anything larger would have resulted in overlapping spheres. Clusters varied substantially in size and some had multiple local maxima (i.e., subclusters). In total, we created 32 spheres across the seven ROIs (7 L dlPFC; 3 R dlPFC; 4 R vlPFC; 7 R dmPFC: 2 L MTG; 5 L SPL; 4 R SPL). Spatial and temporal variability estimates were computed using their respective GLMs (described below) with this set of spheres. Estimates of variability from each sphere were then submitted to multi-level measurement models for further analysis (described in the ‘Statistical Approach’ section). Figure 2 provides an illustration of our ROI definition. Because our focus was on understanding variability in the regions known to instantiate top-down emotion regulation, we did not evaluate either type of variability in the amygdala.

**Figure 2.**
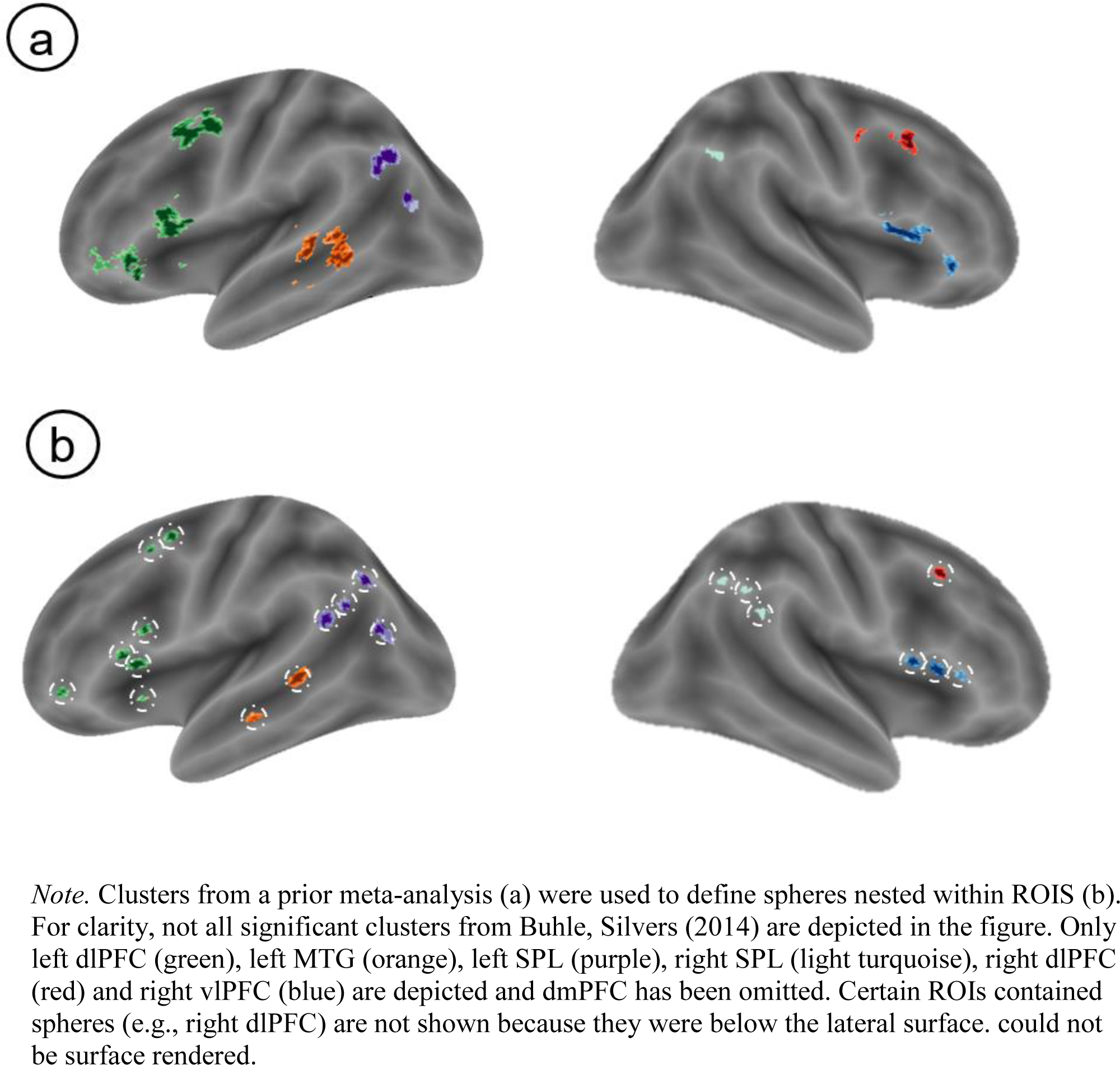
Illustrative Schematic of ROI Definition.

*Spatial Variability Estimation.* We first submitted each participant’s data to a fixed effects analysis using a standard general linear model (GLM) in FSL. The reappraisal task was modeled with five task regressors, each convolved with the canonical HRF (double gamma). A regressor for the instructional cue (‘Instruction’), one for each task condition (‘Far’, ‘Look-Negative’, & ‘Look-Neutral’), and a final regressor for the affect rating period (‘Rating’) were modeled. Slice-timing effects were corrected for by including the temporal derivative of each task regressor in the model. Rotation and translation parameters obtained from MCFLIRT, along with their derivatives and the squares of each, were added as nuisance regressors to reduce the effects of head motion on functional data. Volumes exceeding 0.9 mm in FD were censored at this stage of this analysis using output from FSL Motion Outliers. A second level analysis, which averaged contrast estimates within subject, was carried out using a fixed effects model by forcing the random effects variance to zero. Registration of functional data to the high resolution structural (MPRAGE) was carried out using FSL’s boundary based registration algorithm (Greve & Fischl, 2009). Each participant’s MPRAGE was then non-linearly registered to the MNI152 template image (10mm warp resolution), and the transformation matrix was subsequently used to warp the functional images.

We used univariate activation estimates from the voxels within each sphere to calculate Gini coefficients, a simple but powerful way to quantify spatial variability (Guest & Love, 2017; Leech et al., 2014; Pyatt, 1976). The Gini coefficient was originally developed to study income inequality within specified geographic locations (e.g., cities, countries; Pyatt, 1976). Gini coefficients can range in value from 0 to 1. In the context of income inequality, a Gini coefficient of 0 means that everyone in a given location has exactly the same income; a coefficient of 1 means that one person has all the income and no one else has any. In the context of fMRI data, Gini coefficients can be used to measure the inequality of activation (i.e., BOLD response) among voxels within a given ROI, with greater inequality of activation implying that a smaller subset of voxels account for a larger portion of BOLD signal within the ROI during a given psychological process. In other words, a higher Gini coefficient (i.e., value closer to 1) represents greater inequality of neural activation within a sphere and means that fewer voxels are accounting for more of the activity within the ROI. As this occurs, more activation is peaked around said voxels and less activity is distributed across the rest of the sphere. Larger Gini coefficients indicate that relatively more activation is distributed across a relatively smaller space. Therefore, higher coefficients correspond to less spatial variability across a sphere. In other words, a greater Gini coefficient indicates activity is confined to fewer voxels and is less free to vary across the sphere (see Figure 3 for a visualization). Gini coefficients are useful because they can yield a single numerical index of spatial variability, thus helping to quantify complex theoretical concepts such as focalization. Our procedure for quantifying spatial variability via Gini coefficients follows.

**Figure 3.**
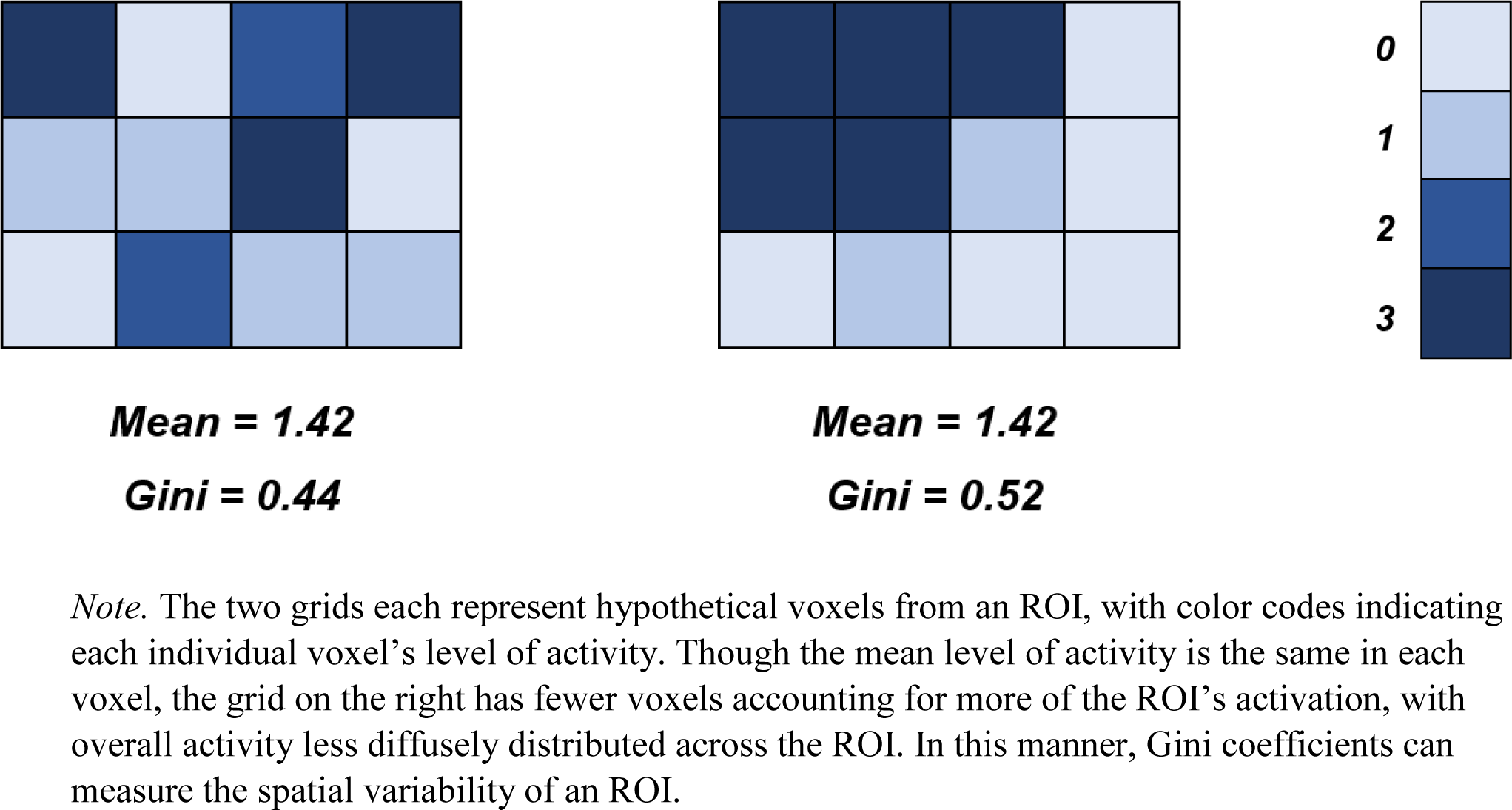
Conceptual Overview of Gini Coefficients and Spatial Variability

As stated previously, our primary focus was on identifying variability during emotion regulation. As such, parameter estimates from the GLM were used to create a linear contrast image comparing the regulation condition (Far) to baseline. These voxelwise contrast values, in the form of Zstat images, were extracted from each sphere using the cosmo_fmri_dataset() command in the MatLab-based CoSMoMVPA toolbox (Oosterhof, Connolly, & Haxby, 2016). The subsequent vector of parameter estimates from each voxel within each sphere was sorted by magnitude and minimum-centered (i.e., we centered at the minimum by subtracting the lowest value in the vector from all elements). Minimum centering ensured that the Gini coefficients remained bound between 0 and 1. Following this step, we used the ordered, minimum-centered vector of parameter estimates to calculate a Gini coefficient. The equation follows.

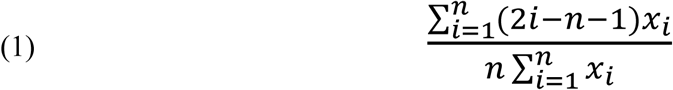

Where *i* represents the rank of a given voxel’s activation, *n* is the total number of voxels, and *xi* is the activation value (i.e., parameter estimate) for the *i*-th voxel. Overall, all subjects had 32 Gini coefficients, each corresponding to a sphere from our ROI list. Importantly, an additional analysis revealed that Gini coefficients were orthogonal to estimates of activity magnitude (details and caveats can be found in the Supplement). Figure 4 displays an overview of Gini coefficient calculation.

**Figure 4.**
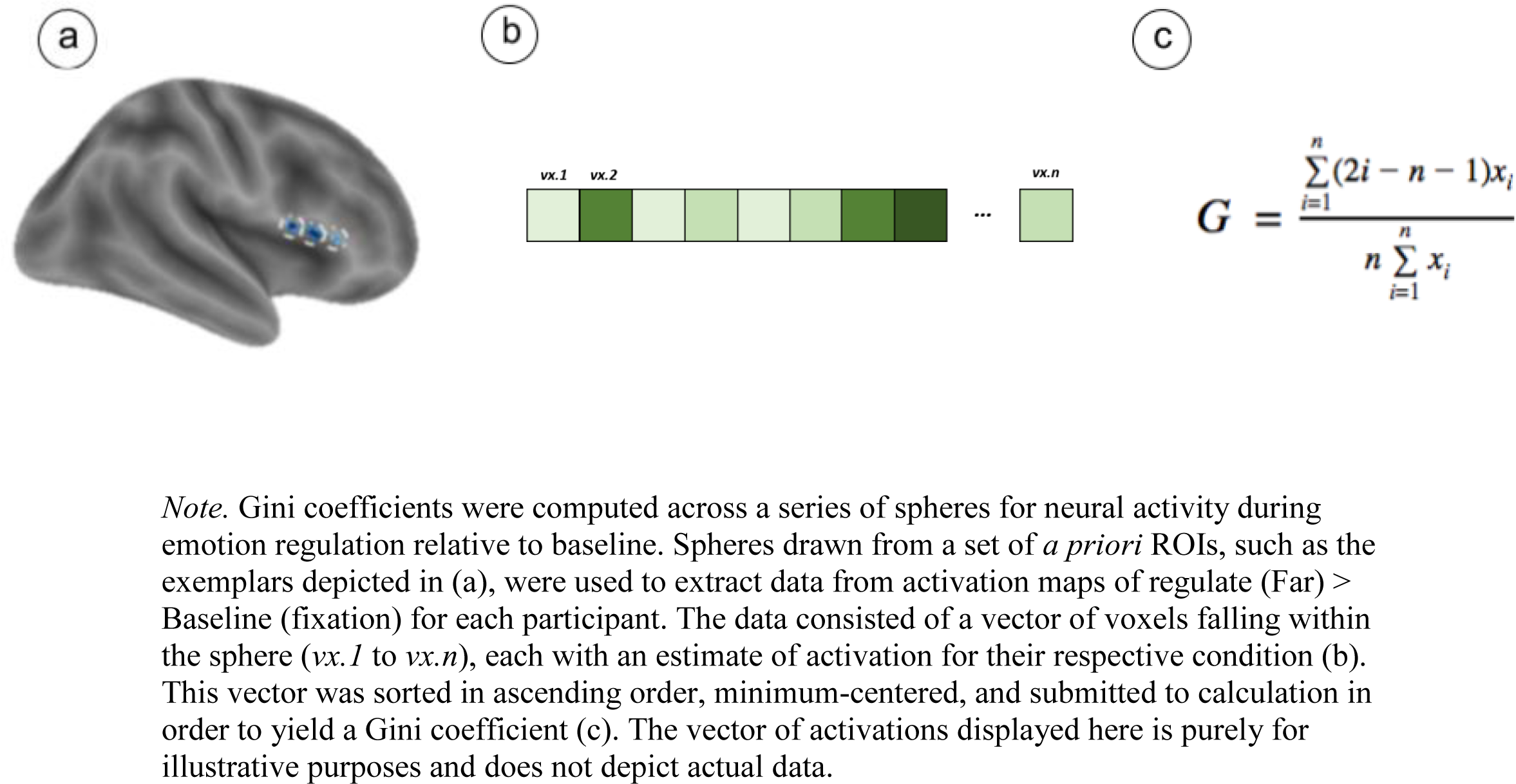
Using Gini Coefficients to Quantify Spatial Variability

*Temporal Variability Estimation*. Temporal variability was estimated by taking the standard deviation of a beta-series (Rissman, Gazzaley, & D’Esposito, 2004) within each sphere. This was executed in three steps. First, we estimated brain activity for each trial using the Least Squares Single (LSS) approach (Mumford, Davis, & Poldrack, 2014; Mumford, Turner, Ashby, & Poldrack, 2012). A fixed-effects GLM was created for each regulation (Far) trial such that the trial of interest was given its own regressor in its own GLM and other trials for that condition were modeled with a separate nuisance regressor. Trials belonging to other conditions (e.g., ‘Look-Negative’, ‘Look-Neutral’, ‘Instruction’, etc.) and motion parameters (standard + extended + volumes exceeding 0.9mm FD) were modeled as their own regressors but not analyzed further. An LSS approach was selected because it is flexible for various types of analyses and results in less multi-collinearity between activation estimates for each trial compared to the Least Squares All (LSA) approach. The second step entailed using parameter estimates from each trial-specific GLM to create a linear contrast image comparing each regulate (Far) trial to baseline. We extracted average estimates of activation within a given sphere across the beta series from the Zmap of the regulate (Far) > baseline contrast. This resulted in an *n* x *p* matrix for each subject, where *n* is the number of trials and *p* is the number of spheres. Each matrix entry represents the mean activity of the *p*-th sphere at the *n*-th trial. The standard deviation of each column was then taken as the third and final step, yielding *p* estimates of temporal variability (where *p* corresponds to the number of spheres) for each subject. Each participant had 32 estimates of temporal variability, each corresponding to a sphere. Figure 5 depicts an overview of this estimation procedure.

**Figure 5.**
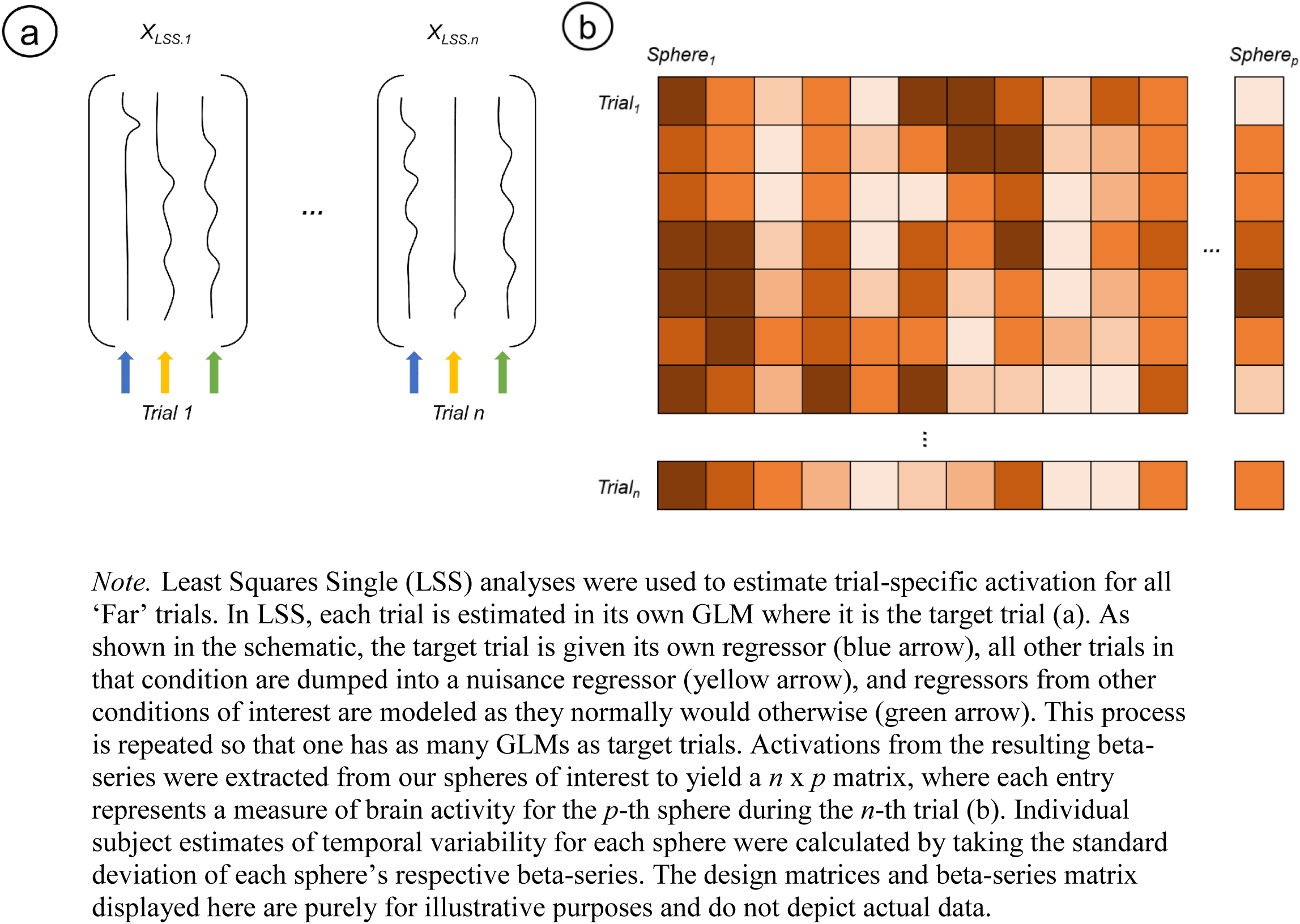
Using Least-Squares Single Trial Analyses to Estimate Temporal Variability.

### Statistical Approach

We used multi-level measurement models to characterize spatial and temporal variability during emotion regulation. These models were further utilized to examine links between variability, age, and affective experience. Below, we describe our motivation for this approach and its benefits before elaborating the modeling procedure in detail. Notably, the variability metrics described above, and the measurement models described below, can be applied to fMRI data obtained with any virtually emotion regulation paradigm1. Moreover, the methods described here can be applied to other conceptualizations of emotion regulation. For instance, one could apply Gini coefficients in conjunction with electroencephalography (EEG) by taking the Gini coefficient of power across different frequency bands, potentially yielding information about whether fewer neural populations are accounting for more of the overall activity. Code and data are publicly available on the open science framework (osf.io/42fkx/).

*Motivation.* As described above, we chose our ROIs based on clusters identified by a recent meta-analysis (Buhle et al., 2014). Importantly, the seven primary clusters of activation reported in this meta-analysis varied widely in terms of voxel size (*k* = 77 to *k* = 517). Using these whole clusters as ROIs would have been problematic, as some clusters would gain enhanced precision from data pooled across a greater number of voxels while making comparisons between regions difficult to interpret. One way to overcome this problem would be to draw a sphere around the global maxima for each cluster. However, this approach is also flawed because it ignores additional information from the rest of the cluster not included in the sphere. The approach used in the present manuscript represent an effort to incorporate as much information as possible from each ROI while maintaining an equal size for each sphere.

The first step of our approach involved drawing spheres (4mm radius) around *all* the maxima (both global and local) within each cluster defined by Buhle et al. (2014). We then estimated spatial and temporal variability within each maxima sphere and importantly, *nested* each local maxima sphere by its ROI (i.e., global maxima). We treated these measurements as manifest (i.e., observed) variables in a multi-level measurement model to estimate latent values for each ROI that varied between subjects. For example, the Buhle et al. meta-analysis identified a 175-voxel cluster in right dlPFC (labeled as right middle frontal gyrus in the meta-analysis). This large cluster’s peak was at X=60, Y=24, Z=3 but also contained two local maxima (X=48, Y=24, Z=9; X=48, Y=15, Z=6). For this particular cluster, three spheres were constructed (one at each peak) and were then nested under a supra-heading of right dlPFC. This allowed us to accurately estimate indices of variability within each ROI that also appropriately incorporated the size of each ROI.

Our multi-level measurement model approach has several benefits. It makes use of data from many spheres within each ROI while using equal sized spheres between ROIs; it minimizes the possibility of including information that was still present in the original ROI-defining meta-analysis due to shared error variance (i.e., from method variance) across studies; it appropriately acknowledges the multi-level structure of the data (multiple measurements nested within participants); it provides a more analytically stable solution than simply running OLS regressions with many terms from our spheres in a single model and avoids the problem of multiple comparisons from running many smaller, ROI-specific OLS regression models.

*Procedure.* After obtaining estimates of spatial and temporal variability, we analyzed these metrics in two separate multi-level measurement models. Specifically, we estimated latent indices of spatial and temporal variability for each ROI and investigated their relationship with age and self-reported affect. Our analytic approach used multi-level modeling to take estimates of variability from spheres drawn around the maxima within each ROI and create a more accurate estimate for the ROI that they comprised. Put another way, the estimates of variability from each sphere of an ROI were used to calculate a latent value for the entire ROI. The relationships between each latent variable (one for each ROI) and its manifest variables (i.e., spheres) were allowed to vary by participant. We further modeled the variance of these latent variables for each ROI as a function of between-person variables of interest: age and average ratings of negative affect during emotion regulation. Two multi-level measurement models—one to measure latent spatial variability and another to measure latent temporal variability—were estimated using Hierarchical Linear Modeling (HLM for windows, version 6.06; Raudenbush & Byrk, 2002).

*Within-Person Spatial Variability Model.* The within-person measurement model for estimating spatial variability follows.

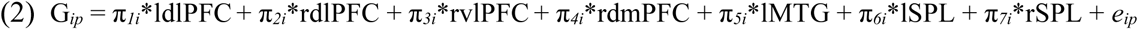

Here, G*ip* represents the Gini coefficient, our index of spatial variability, for the *i*-th individual at the *p*-th sphere. The slopes in this equation (π*1i* – π*7i*) each represent participant-specific, latent Gini coefficients for a given ROI. That is, they are the idealized estimates of Gini coefficients for each ROI, purged of measurement error. Variables labeled by ROI (e.g., ldlPFC, lMTG, etc) reflect overparameterized dummy codes that signify which ROI each G*ip* belonged to. In other words, Equation 2 creates a latent, idealized Gini coefficient for every ROI using the observed Gini coefficients from each ROI’s constituent spheres while discarding measurement error (*eij*). Notably, the lack of an intercept in this model is intentional, as it allows information about average Gini coefficients for each ROI to be encoded in the slopes, (i.e., π’s).

*Within-Person Temporal Variability Model.* The within-person measurement model for estimating temporal variability followed a nearly identical equation as the preceding model.

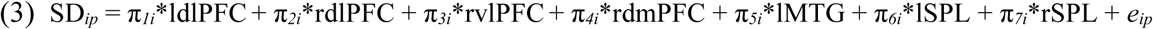

SD*ip* represents the standard deviation of the *p*-th sphere’s beta-series for the *i*-th individual. Paralleling the previous equation, slopes in this equation represent idealized, participant-specific estimates of temporal variability. The same set-of overparameterized dummy codes were used to indicate to which ROI each sphere’s temporal variability estimate “belonged”.

*Between-Person Model.* Between-person equations for the two models were identical in form (but of course differentially represented spatial and temporal variability):

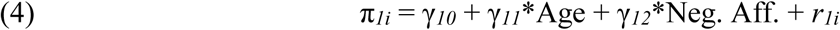

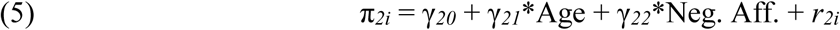

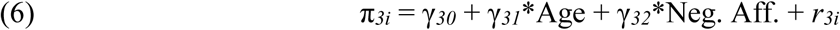

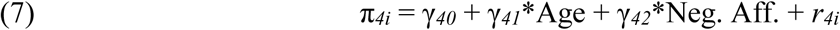

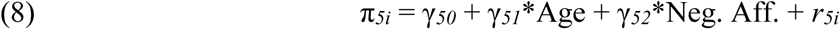

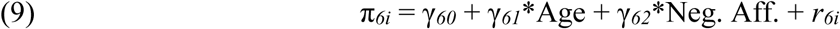

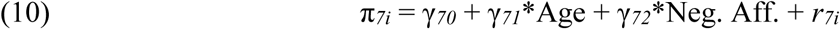

In the between-person part of the model, participant-specific latent variables (π’s) are modeled as a function of overall fixed effects for the latent variables, age, and mean self-reported negative affect during active regulation (γ’s). The addition of participant specific deviations to the between-person part of the model allowed the within-person latent variables to vary between individuals (*r*’s). This allows us to test how age and affective experience relate to participants’ spatial and temporal variability for each ROI. Both age and average ratings of negative affect (Neg. Aff.) were grand mean centered before being entered into the model.

## Results

*Behavioral Results.* The average rating of mean participant negative affect during Far trials was 1.948 (SD = .499), meaning that our participants were on average rating the negative stimuli during reappraisal as inducing low-medium negative affect. The average ratings of negative affect during Look-Negative and Look-Neutral trials were 2.48 (SD = .562) and 1.08 (SD = 0.171), respectively. We conducted a paired samples t-test between average Far and Look-Negative ratings as a manipulation check, revealing significantly different means (*t*(69) = -10.37, *p* < .001). Age was unrelated to mean ratings of negative affect during Far-Negative (*r* = -0.092, *p* > .250), Look-Negative (*r* = 0.184, *p* = .128), and Look-Neutral (*r* = 0.117, *p* > .250) trials. However, the capacity to reappraise—defined as the percent change in ratings of negative affect between the Far and Look-Negative conditions—increased with age (*r* = 0.377, *p* < .01), consistent with prior work (Silvers et al., 2012). As noted earlier, the focus of the study was on emotion regulation so all subsequent analyses focus only on the Far condition.

*Gini Coefficient Validation*. Given that Gini Coefficients have only recently been introduced to neuroimaging research (Guest & Love, 2017; Leech et al., 2014), it was necessary to validate this novel technique in three ways. We first confirmed that Gini coefficients were not simply capturing information about the magnitude of neural activity. That is, our Gini coefficients were orthogonal to ROI means and peaks and thus represent information about *variability* and not magnitude (page 2 of Supplement). Following this, we next tested whether Gini coefficients reflected greater focalization of activation. Consistent with this hypothesis, we found that greater Gini coefficients reflected the spatial coalescence of an ROI’s most active voxels. That is, the most active voxels in a given ROI were more likely to be closer together and not distributed across the entire sphere. Last, we sought to determine whether Gini coefficients were actually simply a proxy for gray matter tissue composition within our ROIs. Results indicated that Gini coefficients are independent of gray matter tissue composition. Statistical output and technical details about these analyses are reported in full in the Supplement.

*Spatial Variability.* As reported in Table 1, relatively similar estimates (ranging from .27-.28) of spatial variability (i.e., Gini coefficients) were observed across ROIs. Low Gini coefficients suggest that activity was distributed somewhat evenly over voxels within each ROI. Given the results of our validation analyses described above (and detailed in the supplement), these Gini coefficients are interpreted as potentially reflecting the presence of a centralized hub of activity, as the most active voxels were concentrated in a focal set of adjacent voxels. The fact that different ROIs had similar mean amounts of spatial variability implies a preserved spatial topography across ROIs during emotion regulation.

**Table 1.**
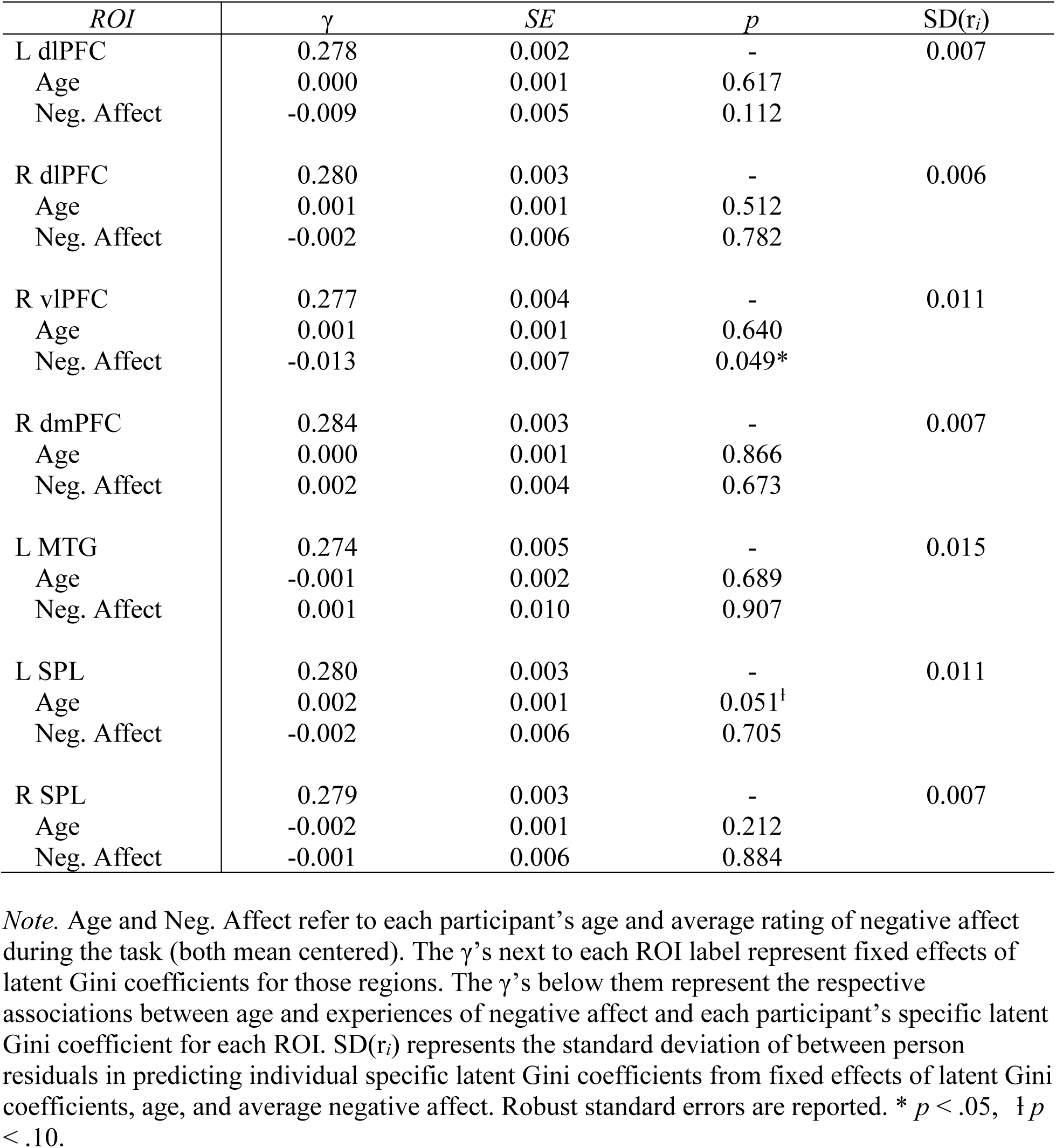
Fixed Effects of Latent Gini Coefficients and Moderators from the Measurement Model

Across participants, estimates of latent spatial variability in left SPL were marginally related to age (γ = .002, SE = .001, *p* = .051; see Table 1). As age increased, fewer voxels in left SPL accounted for more activation than in younger participants. In terms of affect, lower levels of negative affect during reappraisal (i.e., more successful emotion regulation) were associated with a greater Gini coefficient in vlPFC activation (γ = -.013, SE = .007, *p* = .049). This means that successful emotion regulation in youth was associated with having fewer voxels in right vlPFC account for more of the activation.

*Temporal Variability.* Results of the temporal variability measurement model are listed in Table 2. In contrast to what was observed with spatial variability, temporal variability varied markedly across the 7 ROIs. Descriptively, temporal variability followed an apparent spatial gradient, such that prefrontal regions generally showed relatively less variability in activation across time compared to parietal and temporal regions.

**Table 2.** Fixed Effects of Latent Temporal Variability and Moderators from the Measurement Model

| ROI | $\gamma$ | SE | p | SD($r_i$) |
| --- | --- | --- | --- | --- |
| L dlPFC | 0.435 | 0.010 | - | 0.076 |
| Age | -0.002 | 0.003 | 0.452 |  |
| Neg. Affect | 0.031 | 0.019 | 0.107 |  |
| R dlPFC | 0.567 | 0.015 | - | 0.110 |
| Age | -0.006 | 0.006 | 0.280 |  |
| Neg. Affect | -0.010 | 0.026 | 0.690 |  |
| R vlPFC | 0.456 | 0.013 | - | 0.088 |
| Age | -0.009 | 0.005 | 0.092<sup>†</sup> |  |
| Neg. Affect | 0.041 | 0.029 | 0.153 |  |
| R dmPFC | 0.529 | 0.011 | - | 0.080 |
| Age | -0.010 | 0.004 | 0.005* |  |
| Neg. Affect | 0.032 | 0.023 | 0.160 |  |
| L MTG | 0.439 | 0.011 | - | 0.068 |
| Age | -0.007 | 0.004 | 0.079<sup>†</sup> |  |
| Neg. Affect | 0.034 | 0.022 | 0.132 |  |
| L SPL | 0.542 | 0.011 | - | 0.081 |
| Age | 0.001 | 0.004 | 0.801 |  |
| Neg. Affect | 0.028 | 0.024 | 0.243 |  |
| R SPL | 0.534 | 0.013 | - | 0.096 |
| Age | -0.006 | 0.005 | 0.243 |  |
| Neg. Affect | 0.015 | 0.031 | 0.614 |  |

*Note.* Age and Neg. Affect respectively refer to each participant's age and average rating of negative affect during the task (both mean centered). The $\gamma$ 's next to each ROI label represent fixed effects of latent temporal variability for those regions. The $\gamma$ 's below them represent the respective associations between age and experiences of negative affect and each participant's specific latent temporal variability for each ROI. SD( $r_i$ ) represents the standard deviation of between person residuals in predicting individual specific latent temporal variability from fixed effects of latent temporal variability, age, and average negative affect. Robust standard errors are reported. \* $p < .05$ , <sup>†</sup> $p < .10$ .

We observed significant, and marginally significant, associations with age in right dmPFC (γ = -.010, SE = .004, *p* = .005), right vlPFC (γ = -.009, SE = .005, *p* = .092), and left MTG (γ = -.007, SE = .004, *p* = .079). Specifically, as age increased temporal variability across these three ROIs decreased. A supplemental analysis showed that estimates of temporal variability were modestly related to gray matter tissue composition (Supplementary Table 4). We thus re-ran our analyses while controlling for gray matter tissue composition in each sphere. Results indicated that the findings with right dmPFC and left MTG remained significant and marginally significant, respectively; the vlPFC trend did not remain marginally significant. Gray matter composition was directly associated with temporal variability in vlPFC and was inversely related to temporal variability in dmPFC; gray matter composition was unrelated to temporal variability for all other ROIs in addition to also having no relationship with age. Detailed statistical output of analyses concerning gray matter composition can be accessed in the Supplement (Supplementary Tables 5 & 6). These results suggest that age is accompanied by more stable, consistent brain activation in fronto-temporal ROIs implicated in emotion regulation, even when controlling for gray matter composition.

## Discussion

Variability is a critical feature of neural activity that has been previously overlooked in developmental neuroscience studies of emotion regulation. The present study sought to quantify two types of within-individual neural variability during emotion regulation—temporal and spatial—and relate them to age and individual differences in affective experience. We found that age was associated with spatial variability in SPL, while affective experience was associated with spatial variability in vlPFC. By contrast, age was linked with temporal variability in MTG and dmPFC. These results help characterize how spatial and temporal profiles of brain activity give rise to emotion regulatory abilities in youth and inform future work aimed at unpacking the role of variability in affective neurodevelopment.

*Spatial Variability Marks Selective Specialization.* We found that less spatial variability in right vlPFC activation was associated with lower levels of average, self-reported negative affect during emotion regulation. We further observed that age predicted less spatial variability in left SPL activation during emotion regulation. Follow-up analyses revealed that ROIs with less spatial variability tended to have highly active voxels tightly coalesced around a central focal point and less activation elsewhere. These results imply that the development of emotion regulation is supported by a shift towards decreased variability and increased spatial specialization in the prefrontal and parietal cortices. While the cross-sectional nature of the present study design precludes formal testing of this hypothesis, it may motivate future longitudinal work aimed at interrogating this possibility. Overall, our findings imply features about mechanistic processes of emotion regulation that are relevant for a variety of theoretical viewpoints.

The present results enhance our understanding of the roles that vlPFC and SPL play in supporting the development of emotion regulation. Prior work suggests that these two regions are recruited to help meet the working memory and attentional demands of cognitive reappraisal (Buhle et al., 2014; Ochsner et al., 2012). Developmentally, both have also been linked to age-related improvements in cognitive control and emotion regulation (Durston et al., 2006; Silvers et al., 2016). The present findings are of tentative use for existing theories that posit that age-related changes in self-regulation hinge, in part, upon the ability to successfully deploy a burgeoning repertoire of executive functions (see Guassi Moreira & Silvers, 2018 and Calkins & Marcovitch, 2010 for overviews). Our results may help to explain prior work demonstrating that youth use effortful emotion regulation strategies like reappraisal less frequently and less effectively than older individuals (Mischel & Mischel, 1983; Mischel & Baker, 1975; Silvers et al., 2016). While prior work has tied the magnitude of prefrontal and parietal activation to developmental differences in emotion regulation (McRae et al., 2012; Silvers et al., 2015, 2016), here we have done the same with variability of activation in these regions. Although we found initial evidence that decreased spatial variability is associated with more successful emotion regulation (i.e., greater ability), it is important to note that we did not link neural variability to youth’s trait-like tendency to engage in emotion regulation (Silvers & Guassi Moreira, 2017). While this possibility is worth consideration for future research, our data can only fuel speculation on this topic. Regardless, the present findings suggest that prefrontal and parietal specialization is also important to consider, in conjunction with overall activation, when characterizing the neurodevelopment of emotion-regulatory abilities.

Outside of vlPFC and SPL, most other regions (e.g., dlPFC, dmPFC, etc) displayed diffuse spatial topographies during reappraisal and spatial variability did not differ as a function of age or affective experience. This suggests that age and affective experience co-vary with spatial variability in a selective subset of regions. This could indicate that (in youth, at least) certain cognitive features of regulation rely on diffusely distributed patterns of activity, whereas other features are instead dependent on a selective set of modular, specialized clusters of activity. If such is the case, it is interesting to consider why different patterns were observed in vlPFC and SPL than other brain regions. Given that these two brain regions have been particularly strongly linked with the variant of reappraisal employed here (distancing; Ochsner, Silvers, & Buhle, 2012), this could suggest that spatial variability is an indicator of age or individual differences only in brain regions most crucial for performing a given cognitive process. This raises the question of whether distinct or similar patterns of spatial variability might be observed for other psychological processes that change during childhood and adolescence. For example, if youth were asked to complete a theory of mind task (e.g., Spunt & Adolphs, 2014), we might expect age and task performance to be associated with less spatial variability in cortical midline structures like dmPFC that support social cognition (Blakemore & Mills, 2014; Pfeifer & Peake, 2012) but not necessarily in vlPFC and SPL. While testing such possibilities is outside of the scope of the present study, the findings presented here may motivate future research aimed at examining how changes in spatial variability contribute to a range of neurodevelopmental processes in social cognitive and affective domains.

Another notion raised by our results—consistent with a dynamic systems view of development that espouses change over time occurs across a multi-level series of self-organizing systems (Smith & Thelen, 2003)—is that each ROI represents a unique hub in an emotion regulation network, functions optimally at different amounts of within-individual variability, and follows its own developmental trajectory, nested within a broader hierarchical trajectory (e.g., a neural network’s trajectory or the brain’s trajectory). This notion follows from the fact that some of our ROIs were more variable than others and not all showed associations with age. Accordingly, the optimal levels of variability in each hub may be age-dependent, such that more variability is necessary at some ages and not during others. If such a ‘variability-architecture’ were to be confirmed by additional work, it would lend further support to extant neurodevelopmental theories of emotion regulation that posit regulation is comprised of emergent, but ultimately separable, psychological modules (Guassi Moreira & Silvers, 2018). Future research can directly test this by comparing ROI variability between youths and adults to identify the optimal, or normative, level of variability for each ROI, while also examining how such variability changes over the course of development to elucidate whether ROIs are similar or disparate in their maturational trajectories. Testing such possibilities, to which our data cannot currently speak to, encapsulates the next step in a line of research examining how spatial variability characterizes the functional architecture of a range of developmental phenotypes.

*Temporal Variability Reflects Increased Stability with Age.* We found that older individuals, relative to younger ones, showed decreased variability in neural responding over the course of the emotion regulation task. Specifically, age was associated with marginally less variable activation in left MTG, and significantly less variability in dmPFC. Changes in dmPFC variability are noteworthy when considering that mentalizing processes are required to track one’s self-regulation progress during reappraisal and other effortful emotion regulation strategies (Ochsner et al., 2012). Decreased temporal variability in dmPFC may be linked to developmental differences in maintaining goal states and self-monitoring, both of which are needed to guide future action (Kurby & Zacks, 2008; Northoff & Bermpohl, 2004; Richmond & Zacks, 2017). Greater temporal variability in dmPFC at younger ages could reflect an unstable or fluctuating ability to implement emotion regulation (Pfeifer & Peake, 2012). If this is the case, we might hypothesize that as individuals mature, their internal representations of self-regulatory states stabilize and dmPFC variability decreases. Despite being unable to empirically verify the plausibility of this notion with our current data, future work using additional experimental paradigms in the scanner may be able to do so.

Another explanation for the temporal variability results has to do with the social nature of the stimuli used in the present study. Specifically, our age-related findings in dmPFC may be partly explained in terms of ongoing maturation of the social brain (Blakemore & Mills, 2014; Pfeifer & Blakemore, 2012; Pfeifer & Peake, 2012). Social cognitive processes change dramatically during the first two decades of life, especially from late childhood through adolescence (Choudhury, Blakemore, & Charman, 2006; Eisenberg, Spinrad, & Knafo-Noam, 1998; Richardson et al., 2018). Such behavioral development has been linked to structural and functional maturation of cortical midline structures, including dmPFC (Blakemore & Mills, 2014; Mills, Goddings, Clasen, Giedd, & Blakemore, 2014; Mills, Lalonde, Clasen, Giedd, & Blakemore, 2014). It is possible that our age results in dmPFC were observed because the pruning of dmPFC pathways promotes a less variable pattern of BOLD activation across repeated instances of regulation (Blakemore & Mills, 2014; Mills, Lalonde, et al., 2014; Pfeifer & Blakemore, 2012; Somerville et al., 2013; Sowell et al., 2001).

Our findings demonstrating an inverse relationship between age and temporal variability are simultaneously consistent and at odds with the existing literature. On the one hand, some prior research in adult samples suggests that greater temporal variability is an adaptive feature of neural responding because it endows the brain with the ability to easily access different ‘network states’ that may be needed to complete mental tasks (Garrett, Kovacevic, Mcintosh, & Grady, 2010; Garrett, Kovacevic, McIntosh, & Grady, 2011; Petroni et al., 2018). Thus, one might predict that age would predict more flexible (i.e., variable) responses. On the other hand, one could argue that older adolescents may simply need to exert less effort while regulating and display reduced temporal variability across repeated instances of regulation as a result. Indeed, our data appear to support this latter account – older individuals showed less temporal variability on a trial-by-trial basis than younger individuals in certain regions, consistent with prior developmental findings in other domains (Koolschijn et al., 2011; Ordaz et al., 2013). Our findings might also be explained by prior work showing that age-related trajectories of temporal variability in the brain do not follow a single pattern, but instead show regionally-specific increases and decreases in variability (Nomi et al., 2017; Petroni et al., 2018). This supports the notion that development is possibly characterized by a process of ‘variability tuning’, where an intermediate amount of variability is necessary for the execution of psychological tasks, but what is classified as “intermediate” varies across brain regions. Our own findings cannot directly confirm or falsify this possibility, but the notion is supported by recently published work which espouses a ‘goldilocks’ view of variability (insufficient or excessive variability is detrimental) (Dinstein, Heeger, & Behrmann, 2015).

*Generalizing to Other Emotion Regulation Strategies.* Though we used fMRI data in conjunction with a cognitive reappraisal task, one strength of the methods and framework introduced here is that they can be applied across a range of emotion regulation paradigms. As we mentioned earlier, humans have an entire toolbox of emotion regulation strategies at their disposal (Braunstein et al., 2017), and cognitive reappraisal is but one strategy. Children and adolescents are able to use other strategies such as distraction and suppression, in addition to ‘implicit’ emotion regulation skills (e.g., reversal learning, extinction) (Guassi Moreira & Silvers, 2018). Future work ought to directly compare developmental trajectories related to neural variability across multiple emotion regulation strategies in order to test whether the present results are unique to reappraisal or generalize to other strategies. For example, one could examine whether neural estimates of spatial and temporal variability change with age and affective experience during emotional suppression in similar or different ways as they do during reappraisal (Goldin, McRae, Ramel, & Gross, 2008; Phan et al., 2005).

*Limitations and Future Directions.* This study is not without limitations, especially since it is the first to examine neural variability in the context of emotion regulation development. First and foremost, since we relied on a cross-sectional design, we cannot characterize true neurodevelopmental trajectories without longitudinal data (e.g., Snijders & Bosker, 1998; Singer & Willett, 2003). Relatedly, without a group of younger children and adults, it is hard to make inferences about developmental processes that extend prior to late childhood and beyond the adolescent years (e.g., a developmental switch; Casey, 2015; Gee et al., 2014) Another limitation lies in our use of measurement models, which provide a simple way to summarize our indices of variability across each ROI, but (possibly incorrectly) assume that the microarchitecture of different cortical spheres is similar in nature.

The ROIs used in the present study were drawn from a meta-analysis that was nearly exclusively based on adult studies. This methodological choice was due to the fact that no meta-analyses based exclusively on pediatric studies exist (Guassi Moreira & Silvers, 2018), and that doing so provided us with a way to independently define ROIs (Kriegeskorte, Simmons, Bellgowan, & Baker, 2009). Unfortunately, this means that the ROIs studied here may not be representative of the precise ROIs that youth use when engaging in emotion regulation. Indeed, it is possible that children use different brain regions than adolescents or adults in order to accomplish the same emotion regulation goals, although it is worth noting this is unlikely given that no prior data to our knowledge have consistently observed activations in children in brain regions not observed in adults. Future work in larger samples might opt to sidestep this issue by defining ROIs in a subset of their sample and using the rest of the sample for independent analyses or by collecting additional runs of an emotion regulation task specifically devoted to ROI localization and identification.

An additional, infrequently discussed limitation revolves around the notion that the scanner is anxiety-provoking and thus constitutes a unique affective context (Eatough, Shirtcliff, Hanson, & Pollak, 2009; Galván, Van Leijenhorst, & McGlennen, 2012). If this were the case, it could impact regulatory processes in ways that obfuscate the true, ecological nature of youth’s emotion regulation. That said, care was taken to reduce this possibility in the present study by familiarizing participants with the scanning environment through the use of mock scanning.

Given that our study was the first to examine variability in youth’s neural responses during emotion regulation, we opted for an fMRI paradigm that was highly consistent with those previously used in youth (McLaughlin, Peverill, et al., 2015; McRae et al., 2012; Silvers et al., 2015). This decision led us to exclusively examine regulation of negative emotion (inclusive of multiple negative emotional states). It is possible that our results might not generalize to regulation of other emotional states and categories. Indeed, other work has suggested that age-related differences in the behavioral and neural correlates of emotion regulation vary across different types of emotional stimuli (Silvers et al., 2012, 2014). Future work ought to examine whether the results reported herein not only generalize across different variants of reappraisal and related emotion regulation strategies, but also across different emotional states.

Last, our study was exploratory by definition because it was the first of its kind. The present results do not confirm any hypotheses, but rather generate novel ones to be tested in future studies. This limitation, like the others above, must be taken into careful consideration when evaluating the impact of our findings with respect to the broader literature.

*Concluding Remarks.* Variability is a fundamental feature of neurodevelopment. Our results suggest that variability is also central to the acquisition of effective emotion regulation. This investigation was the first to incorporate neural variability into developmental research on emotion regulation. We showed that age was associated with increasing spatial focalization of activity in some brain regions, whereas other regions exhibited spatial topographies that were age- and experience-invariant. We also showed that age is marked by stability of neural responses over the course of repeated emotion regulation. Overall, this work contributes towards identifying the mechanisms that encode the neural bases of emotion regulation in youth.

## Acknowledgements

This research was supported by a grant (#MH103291) from the National Institute of Mental Health awarded to KAM. Preparation of this manuscript was supported by a National Science Foundation Graduate Research Fellowship (Fellow ID: 2016220797) to JFGM. Gratitude is extended towards Drs. Carolyn Parkinson and Jesse Rissman for their feedback about the study concept and towards Emilia Ninova for reflections on the manuscript.

## Supplement

### Univariate fMRI Data Pre-Processing, Analysis, and Results

*Pre-Processing.* Prior to pre-processing, data were visually inspected for artifacts and anatomical abnormalities. fMRI data were pre-processed and analyzed using the fMRI Expert Analysis Tool (FEAT, version 6.00) of the FMRIB Software Library package (FSL, version 5.0.9; fsl.fmrib.ox.ac.uk). Pre-processing consisted of using the brain extraction tool (BET) to remove non-brain tissue from functional and structural runs, spatial realignment of functional volumes to correct for head motion using MCFLIRT, and hi-pass filtering the data (100s cutoff). The extent of participant head motion was further estimated by running FSL Motion Outliers to record volumes that exceeded a 0.9 mm threshold of framewise displacement (FD) (Siegel et al., 2014). Runs were discarded if participants exceeded this threshold for more than 25 volumes (~20% of a single run). Functional data were smoothed using a 5 mm Gaussian kernel, full-width-at-half maximum to increase signal-to-noise ratio. Data were pre-whitened to account for serial autocorrelation across time.

*First-Level Analysis.* We first submitted each participant’s data to a fixed effects analysis in FSL using the general linear model (GLM). The reappraisal task was modeled with five task regressors (specified with FSL’s three-column format), each convolved with the canonical HRF (double gamma). A regressor for the instructional cue (‘Instruction’), one for each task condition (‘Far’, ‘Look-Negative’, & ‘Look-Neutral’), and a final regressor for the affect rating period (‘Rating’) were modeled. Slice-timing effects were corrected for by including the temporal derivative of each task regressor in the model. Rotation and translation parameters obtained from MCFLIRT, along with their derivatives and the squares of each, were added as nuisance regressors to reduce the effects of head motion on functional data. Volumes exceeding 0.9 mm in FD were also censored at this stage of this analysis using output from FSL Motion Outliers. Each participant’s functional runs were averaged together and then registered to their MPRAGE scan. Each participant’s MPRAGE was non-linearly registered to the MNI152 template image (10mm warp resolution), and the transformation matrix was subsequently used to warp the functional images.

*Group-Level Analysis for Regulation > Uninstructed Viewing (Aversive).* The parameter estimates from this GLM were then used to create linear contrast images comparing conditions of interest. We created a linear contrast comparing active regulation (‘Far’) to uninstructed viewing of aversive stimuli (‘Look-Negative’). Random effects, group-level analyses were performed on this contrast using FSL. Thresholded Z-statistic images were computed to examine overall differences in neural activation for these two contrasts of interest using FSL’s FLAME1 module (Beckmann et al., 2003). An additional whole-brain analysis was conducted in which participant ages, centered at the mean, were regressed on the regulate (‘Far’) > uninstructed viewing (aversive, ‘Look-Neg’), to understand the relationship between age and magnitude of neural activation during regulation. All thresholded images were obtained with a cluster defining threshold of 3.1 and a familywise error rate of *p* < 0.05 (Random Field Theory cluster correction; Poline et al., 1997). Results depicted in Supplementary Figure 1 are projected onto an average of all participants’ structural images (i.e., the bg_image.nii.gz file found in the higher level FEAT directory).

*Results.* As illustrated in Table 1 and Supplementary Figure 1 (Panel A), participants exhibited greater activity in right motor cortex when regulating affective responses to aversive images compared to uninstructed viewing. Additionally, participants recruited the left temporo-parietal junction (TPJ) (Figure 1, Panel B). By contrast, uninstructed viewing of negative stimuli was linked with greater recruitment of several early visual processing areas as well as activation in the motor cortex, albeit on the opposite side (left; not pictured). No significant clusters survived correction when regressing age on this contrast.

**Supplementary Figure 1.**
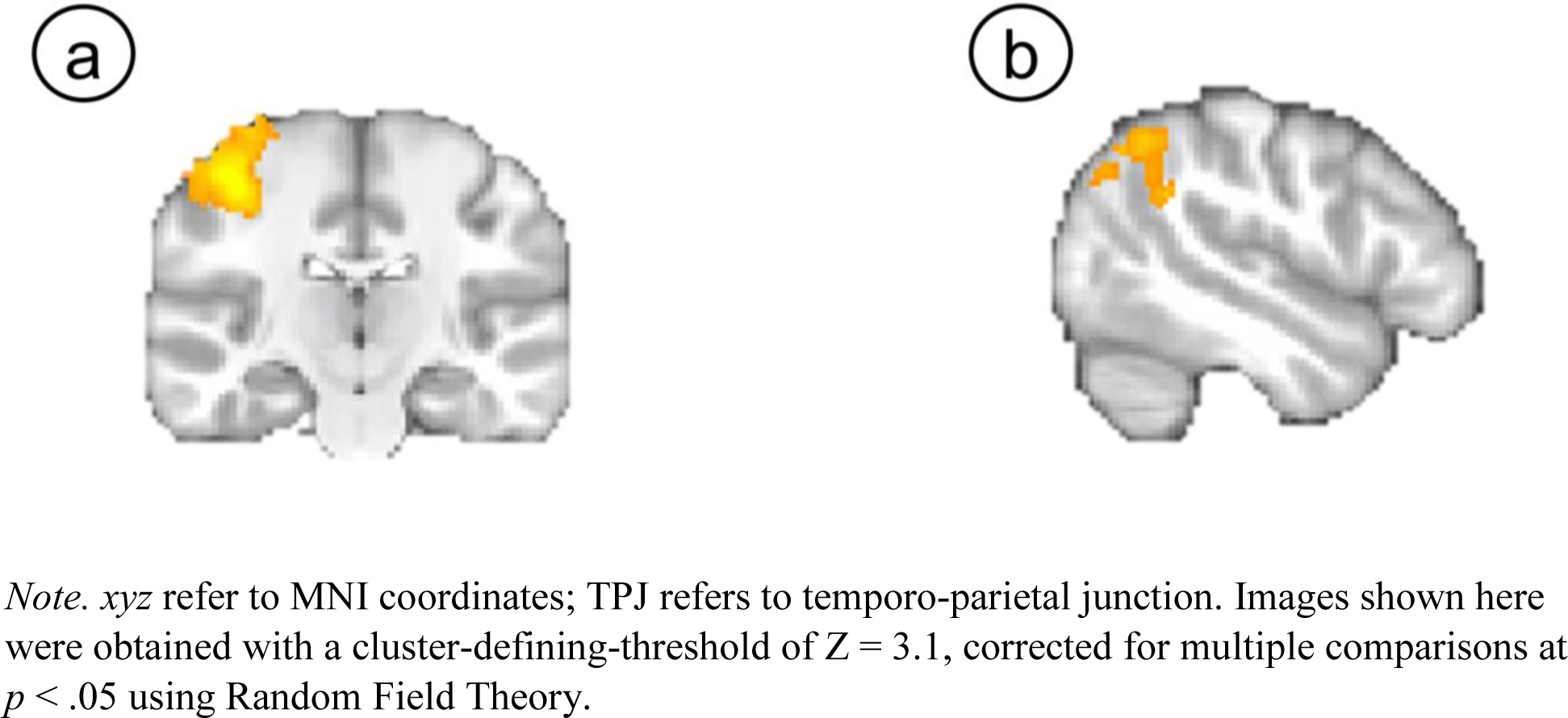
Activation for Regulate (‘Far’) > Uninstructed Viewing (Aversive; ‘Look-Negative’). We found a broad cluster of activation in right motor cortex (*xyz* = 42, -20, 54) (a) in addition to activation in a cluster encompassing the left TPJ (*xyz* = -48, -56, 52) (b).

*Note. xyz* refer to MNI coordinates; TPJ refers to temporo-parietal junction. Images shown here were obtained with a cluster-defining-threshold of Z = 3.1, corrected for multiple comparisons at *p* < .05 using Random Field Theory.

### Distinctions Between Variability and Magnitude Estimates of Brain Activation

Given the paucity of neuroimaging research using Gini coefficients, we investigated whether Gini coefficients captured distinct information from standard univariate analyses of mean or peak activation. To do so, we extracted peak and mean estimates of activation from the Regulate (Far) > baseline contrast using spheres constructed around the global maxima of each ROI in our set. Since this analysis only examined one maxima (global) of each ROI from the Silvers, Buhle et al meta-analysis (2014), we broadened our sphere radius to 6 mm2. This resulted in three metrics per each of the seven ROIs: peaks, means, and Gini coefficients. Next, we correlated these three metrics to determine whether the information encoded by Gini coefficients was orthogonal to mean and peak estimates of signal intensity. Across all triplets of Gini, mean, and peak estimates, the only significant correlation between a Gini coefficient and an estimate of magnitude was in right dlPFC. In dlPFC (R) Gini coefficients were observed to be significantly correlated with peak estimates (*r* = 0.299, *p* = .012; all correlations displayed in Supplementary Table 2). However, this value did not survive correction for multiple comparisons a la Holm (corrected *p*-value threshold = 0.004). Estimates of temporal variability were not examined since prior work has previously found them to be orthogonal to estimates of magnitude (Garrett, Kovacevic, McIntosh, & Grady, 2011). Therefore, we conclude that the Gini coefficients in our data encoded unique information about neural activity.

**Supplementary Table 1.** Brain regions which showed significant activation during emotion regulation compared to uninstructed viewing of aversive images (‘Far’ > ‘Look-Neg’).

| *Region* |  | *x* | *y* | *z* | *Z* | *k* |
| --- | --- | --- | --- | --- | --- | --- |
| *Far > Look-Negative* |  |  |  |  |  |  |
| Primary Somatosensory Cortex | R | 44 | -22 | 56 | 5.6 | 1204<sup>a</sup> |
| Primary Motor Cortex | R | 38 | -24 | 56 | 5.49 | <sup>a</sup> |
| Pre-Motor Cortex | R | 36 | -16 | 70 | 4.94 | <sup>a</sup> |
| Primary Somatosensory Cortex | R | 52 | -16 | 52 | 4.57 | <sup>a</sup> |
| Pre-Motor Cortex | R | 32 | -26 | 74 | 3.73 | <sup>a</sup> |
| TPJ | L | -52 | -50 | 32 | 4.56 | 466<sup>b</sup> |
| IPL | L | -52 | -54 | 50 | 4.49 | <sup>b</sup> |
| IPL | L | -52 | -56 | 34 | 4.28 | <sup>b</sup> |
| IPL | L | -50 | -60 | 44 | 4.27 | <sup>b</sup> |
| IPL | L | -50 | -64 | 40 | 4.17 | <sup>b</sup> |
| IPL | L | -48 | -64 | 46 | 3.89 | <sup>b</sup> |
| *Look-Negative > Far* |  |  |  |  |  |  |
| Visual Cortex (V3) | L | -10 | -82 | -6 | 7.89 | 5599<sup>c</sup> |
| Visual Cortex (V1) | R | 10 | -84 | 6 | 6.56 | <sup>c</sup> |
| Visual Cortex (V2) | R | 12 | -76 | -8 | 6.39 | <sup>c</sup> |
| Visual Cortex (V1) | R | 18 | -84 | 10 | 6.35 | <sup>c</sup> |
| Visual Cortex (V4) | R | 24 | -70 | -12 | 6.12 | <sup>c</sup> |
| LOC | R | 20 | -84 | 32 | 5.55 | <sup>c</sup> |
| IPL | L | -60 | -22 | 38 | 6.32 | 4502<sup>d</sup> |
| IPL | L | -60 | -26 | 26 | 6.29 | <sup>d</sup> |
| Primary Somatosensory Cortex | L | -50 | -22 | 54 | 6.23 | <sup>d</sup> |
| SPL | L | -34 | -52 | 62 | 5.75 | <sup>d</sup> |
| SPL | L | -32 | -46 | 64 | 5.53 | <sup>d</sup> |
| Primary Somatosensory Cortex | L | -32 | -32 | 58 | 5.53 | <sup>d</sup> |
| LOC | L | -24 | -80 | 28 | 4.38 | 221<sup>e</sup> |
| aIPS | L | -28 | -68 | 30 | 3.97 | <sup>e</sup> |
| LOC | L | -24 | -86 | 18 | 3.59 | <sup>e</sup> |

*Note.* R = right and L = left; x, y, and z refer to MNI coordinates. Regions that share the same superscript are part of the same cluster. 'Far' refers to active regulation; 'Look-Negative' refers to uninstructed viewing of negative stimuli; 'Look-Neutral' refers to uninstructed viewing of neutral stimuli; TPJ refers to temporo-parietal junction; IPL refers to inferior parietal lobule; LOC refers to lateral occipital cortex; SPL refers to Superior Parietal Lobule; aIPS refers to anterior intra-parietal sulcus.

**Supplementary Table 2.** Correlations between Gini coefficients (larger Gini coefficients interpreted as less spatial variability) and estimates of magnitude (mean, peak) for spheres corresponding to the global maxima of each ROI.

| *ROI* | *r(Gini, Peak), p* | *r(Gini, Mean), p* |
| --- | --- | --- |
| L dlPFC | 0.212, 0.078 | 0.096, 0.431 |
| R dlPFC | 0.299, 0.012 | 0.013, 0.915 |
| R vlPFC | 0.172, 0.155 | -0.026, 0.829 |
| R dmPFC | -0.002, 0.986 | -0.071, 0.561 |
| L MTG | 0.087, 0.475 | -0.210, 0.080 |
| L SPL | 0.168, 0.163 | -0.174, 0.149 |
| R SPL | 0.128, 0.293 | -0.014, 0.905 |

*Note.* The first entry in each cell represents a correlation coefficient between Gini coefficients and an estimate of magnitude (peaks in the first column, means in the second). The second entry is an uncorrected p-value. None of the p-values displayed here survived multiple comparison correction. This analysis demonstrates that Ginis encode information that is orthogonal to estimates of magnitude (mean and peak).

### Gini Coefficients Potentially Reflect the Presence of Centralized Nuclei

We argue that larger Gini coefficients represent decreased spatial variability in the main document. However, this diminished spatial variability could manifest itself in two ways. One possibility is that activation has become consolidated into a single, central nucleus somewhere within an ROI. Another possibility is that a higher Gini coefficient represents the coalescence of activity into a set of smaller, distributed nodes that form a micro-network within the ROI. While both cases potentially reflect the consolidation of information in a specialized set of voxels, it was unknown whether this occurred in a centralized or distributed fashion. We evaluated these possibilities by first using CoSMoMVPA to extract activity estimates from the global maxima of each ROI in our set (6mm radius sphere) and discarded any voxels outside the top 10% in terms of activity. Afterwards, we computed the standard deviation of the x, y, and z coordinates of these voxels and then took the average of the three standard deviations. This index describes the manner in which the most active voxels within the sphere were distributed in relation to each other: a lower number indicates the most active voxels were relatively close to another, whereas a higher number indicates the most active voxels were relatively far away. We then correlated these estimates with Gini coefficients for each ROI across participants. Summarized in Supplementary Table 3, results showed that having a larger Gini coefficient was related to having the most active voxels relatively more adjacent in space (*r* coefficients were all negative and ranged between -0.142 and -0.479). These supplementary results indicate that greater Gini coefficients imply the presence of a centralized nucleus within ROIs.

**Supplementary Table 3.**
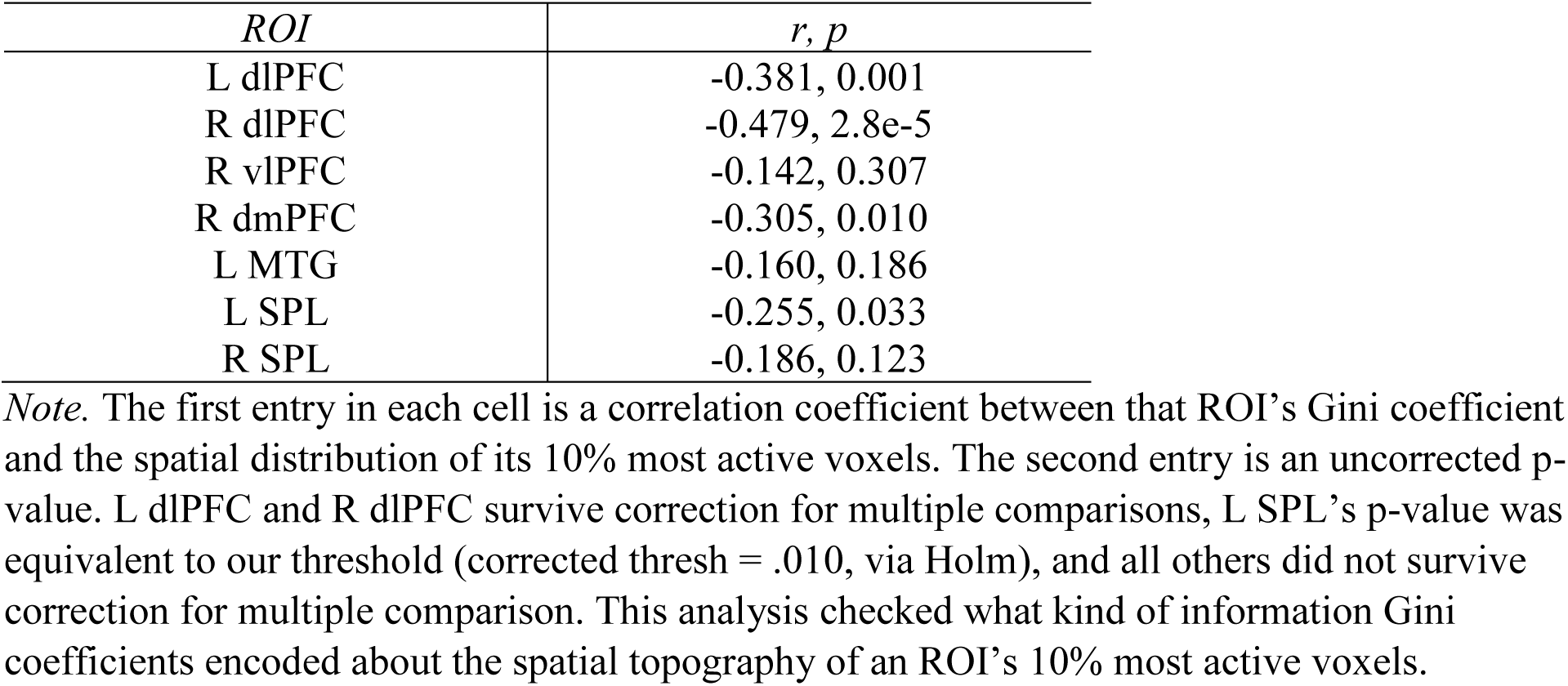
Correlations between Gini coefficients and proximity of the 10% most active voxels within each ROI.

| *ROI* | *r, p* |
| --- | --- |
| L dlPFC | -0.381, 0.001 |
| R dlPFC | -0.479, 2.8e-5 |
| R vlPFC | -0.142, 0.307 |
| R dmPFC | -0.305, 0.010 |
| L MTG | -0.160, 0.186 |
| L SPL | -0.255, 0.033 |
| R SPL | -0.186, 0.123 |

*Note.* The first entry in each cell is a correlation coefficient between that ROI's Gini coefficient and the spatial distribution of its 10% most active voxels. The second entry is an uncorrected p-value. L dlPFC and R dlPFC survive correction for multiple comparisons, L SPL's p-value was equivalent to our threshold (corrected thresh = .010, via Holm), and all others did not survive correction for multiple comparison. This analysis checked what kind of information Gini coefficients encoded about the spatial topography of an ROI's 10% most active voxels.

### Controlling for Gray Matter Composition

An important concern regarding our estimation of variability lies in the role of gray matter composition. Since prior research has indicated that gray matter changes drastically across juvenile development (Mills, Lalonde, Clasen, Giedd, & Blakemore, 2014; Sowell, Thompson, Tessner, & Toga, 2001), it possible that developmental differences in gray matter could have confounded our results. To examine this issue, we first tested to see whether estimates of each ROI’s gray matter composition (compared to white matter and cerebrospinal fluid) were related to our estimates of variability and then, if necessary, subsequently re-ran our measurement models controlling for gray matter.

To address this issue, we first set about estimating voxelwise tissue composition among our 32 spheres for each participant. This was accomplished using FMRIB’s Automated Segmentation Tool (FAST), which segments structural brain images into gray matter, white matter, and cerebrospinal fluid using an expectation-maximization algorithm in conjunction with hidden Markov random field models (Zhang, Brady, & Smith, 2001). The code used to run FAST in the current study was: fast -t 1 -n 3 -H 0.1 -I 4 -l 20.0 -g -o t1_struct_image.nii.gz (a full script with our implementation is available on github via the OSF).

The resulting output describes the extent to which each voxel comprises gray matter, white matter, and cerebrospinal fluid (as a percentage). Afterwards, each participant’s gray matter composition image (i.e., pve_1.nii.gz) was masked with all 32 spheres and gray matter composition values were extracted from each. The average values for each ROI are depicted in Supplementary Table 4: the majority of all ROIs were composed of gray matter.

**Supplementary Table 4.** Average gray matter composition of each ROI.

| *ROI* | *Mean (SD)* |
| --- | --- |
| L dlPFC | 0.547 (0.239) |
| R dlPFC | 0.545 (0.191) |
| R vlPFC | 0.556 (0.256) |
| R dmPFC | 0.643 (0.229) |
| L MTG | 0.553 (0.200) |
| L SPL | 0.633 (0.286) |
| R SPL | 0.762 (0.217) |

*Note.* Composition values expressed as percentages such that each value reflects the percentage of gray matter (compared to white matter and cerebrospinal fluid) constituting each ROI. Values were obtained from FSL's FAST segmentation algorithm and were averaged across participants and multiple spheres for each ROI. *SD* refers to standard deviation.

Following this, we tested the extent to which gray matter composition might be statistically related to our estimates of variability. To this end, we ran two random coefficient regressions of the following single-equation form.

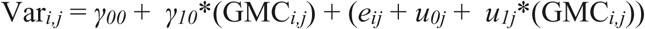

Where the *i*th variance estimate from a given sphere for the *j*th participant is modeled as a function of the corresponding gray matter composition (GMC) from that participant’s sphere. Two models were run: one with spatial variability estimates and another with temporal variability estimates. Parameter estimates represent the expected variance estimate when gray composition is zero (*γ00*) and the linear dependency between gray matter composition and variability (*γ10*). Crucially, this analysis adequately handles the multi-level nature of the data (repeated measurements nested within participants) and creates non-biased and efficient standard errors with which to test parameter estimates with. Results showed that gray matter composition was unrelated to estimates of spatial variability (*γ10* = 0.001, SE = 0.005, *t* = 0.17, *ns*), but was directly related with estimates of temporal variability (*γ10* = 0.0483, SE = 0.012, *t* = 4.02, *p* < .001). In light of these results, we re-ran our multi-level measurement models while controlling for the average gray matter composition of each ROI. Notably, our significant finding with dmPFC temporal variability and age remained significant (see Supplementary Table 5). We are thus left to conclude that available indicates our results are robust to developmental differences in gray matter composition.

**Supplementary Table 5.**
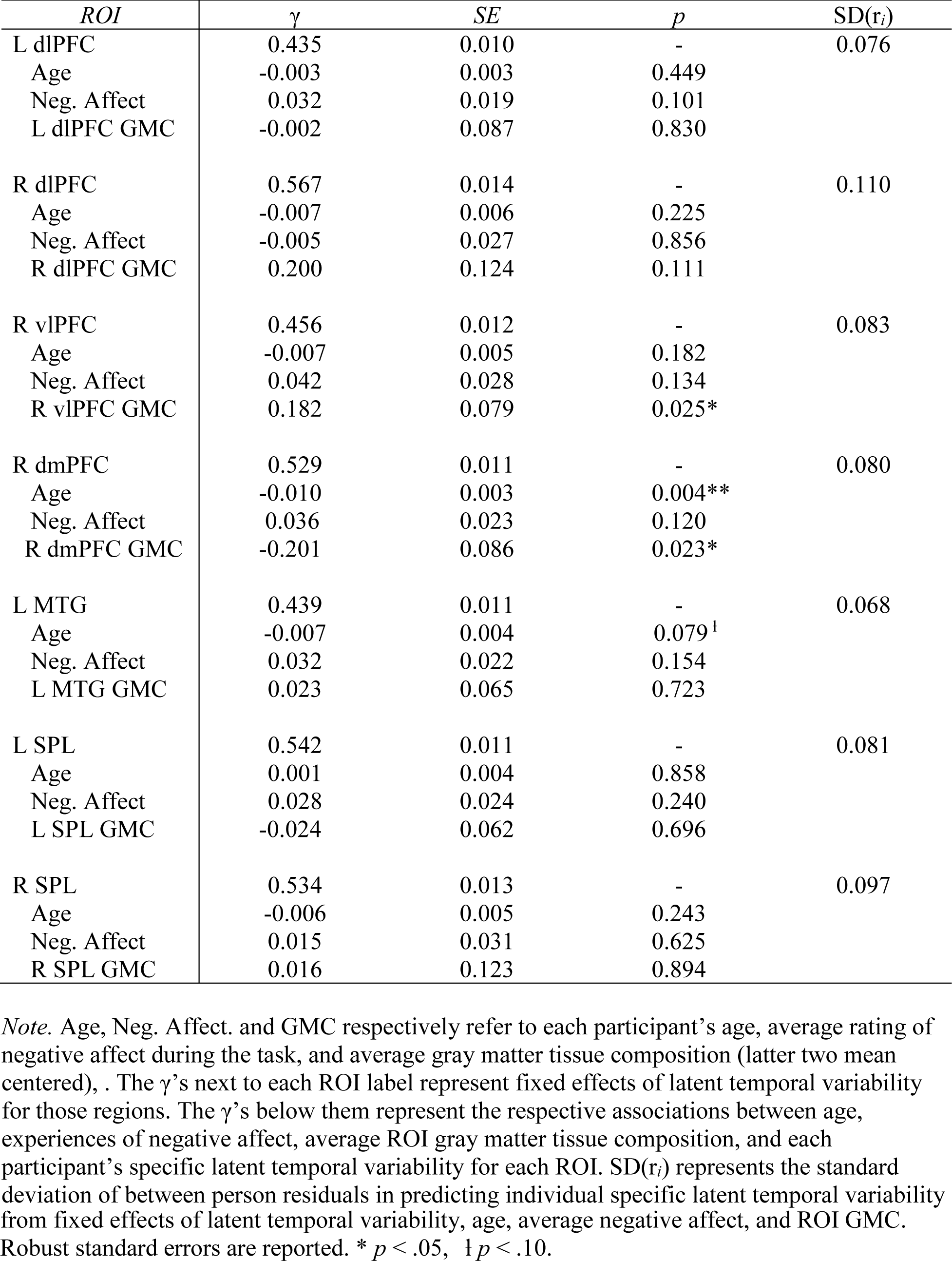
Fixed Effects of Latent Temporal Variability and Moderators from the Measurement Model, Controlling for Average ROI Gray Matter Composition

| *ROI* | $\gamma$ | *SE* | *p* | *SD(r<sub>i</sub>)* |
| --- | --- | --- | --- | --- |
| L dlPFC | 0.435 | 0.010 | - | 0.076 |
| Age | -0.003 | 0.003 | 0.449 |  |
| Neg. Affect | 0.032 | 0.019 | 0.101 |  |
| L dlPFC GMC | -0.002 | 0.087 | 0.830 |  |
| R dlPFC | 0.567 | 0.014 | - | 0.110 |
| Age | -0.007 | 0.006 | 0.225 |  |
| Neg. Affect | -0.005 | 0.027 | 0.856 |  |
| R dlPFC GMC | 0.200 | 0.124 | 0.111 |  |
| R vlPFC | 0.456 | 0.012 | - | 0.083 |
| Age | -0.007 | 0.005 | 0.182 |  |
| Neg. Affect | 0.042 | 0.028 | 0.134 |  |
| R vlPFC GMC | 0.182 | 0.079 | 0.025* |  |
| R dmPFC | 0.529 | 0.011 | - | 0.080 |
| Age | -0.010 | 0.003 | 0.004** |  |
| Neg. Affect | 0.036 | 0.023 | 0.120 |  |
| R dmPFC GMC | -0.201 | 0.086 | 0.023* |  |
| L MTG | 0.439 | 0.011 | - | 0.068 |
| Age | -0.007 | 0.004 | 0.079<sup>†</sup> |  |
| Neg. Affect | 0.032 | 0.022 | 0.154 |  |
| L MTG GMC | 0.023 | 0.065 | 0.723 |  |
| L SPL | 0.542 | 0.011 | - | 0.081 |
| Age | 0.001 | 0.004 | 0.858 |  |
| Neg. Affect | 0.028 | 0.024 | 0.240 |  |
| L SPL GMC | -0.024 | 0.062 | 0.696 |  |
| R SPL | 0.534 | 0.013 | - | 0.097 |
| Age | -0.006 | 0.005 | 0.243 |  |
| Neg. Affect | 0.015 | 0.031 | 0.625 |  |
| R SPL GMC | 0.016 | 0.123 | 0.894 |  |

*Note.* Age, Neg. Affect. and GMC respectively refer to each participant's age, average rating of negative affect during the task, and average gray matter tissue composition (latter two mean centered), . The $\gamma$ 's next to each ROI label represent fixed effects of latent temporal variability for those regions. The $\gamma$ 's below them represent the respective associations between age, experiences of negative affect, average ROI gray matter tissue composition, and each participant's specific latent temporal variability for each ROI. *SD(r<sub>i</sub>)* represents the standard deviation of between person residuals in predicting individual specific latent temporal variability

**Supplementary Table 6.** Correlations between average gray matter composition for each ROI and participant age.

| *ROI* | *r, p* |
| --- | --- |
| L dlPFC | -0.058, 0.636 |
| R dlPFC | 0.095, 0.436 |
| R vlPFC | -0.202, 0.094 |
| R dmPFC | 0.019, 0.876 |
| L MTG | -0.143, 0.236 |
| L SPL | -0.252, 0.036 |
| R SPL | -0.021, 0.861 |

*Note.* The first entry in each row is a correlation coefficient between that ROI's mean gray matter composition values and participant age. The second entry is an uncorrected $p$ -value. Composition values were calculated as percentages such that each value reflects the percentage of gray matter (compared to white matter and cerebrospinal fluid) constituting each ROI. Values were obtained from FSL's FAST segmentation algorithm. No values survive correction for multiple comparison ( $p = .007$ threshold).

## Footnotes

1 Some kind of beta-series or timeseries is required for estimation of temporal variability, but most fMRI tasks can be marshalled to fit this prerequisite. Multiple spheres nested within multiple ROIs are required for the multilevel measurement models.

2 This could not be done for analyses in the main document, as spheres would have then substantially overlapped.

## References

Aldao, A., Sheppes, G., & Gross, J. J. (2015). Emotion Regulation Flexibility. Cognitive Therapy and Research, 39(3), 263–278. https://doi.org/10.1007/s10608-014-9662-4

Blakemore, S.-J., & Mills, K. L. (2014). Is adolescence a sensitive period for sociocultural processing? Annual Review of Psychology, 65, 187–207. https://doi.org/10.1146/annurev-psych-010213-115202

Braunstein, L. M., Gross, J. J., & Ochsner, K. N. (2017). Explicit and implicit emotion regulation: A multi-level framework. Social Cognitive and Affective Neuroscience, *ePub ahead*, 1–13. https://doi.org/10.1080/02699931.2010.544160

Buhle, J. T., Silvers, J. A., Wage, T. D., Lopez, R., Onyemekwu, C., Kober, H., … Ochsner, K. N. (2014). Cognitive reappraisal of emotion: A meta-analysis of human neuroimaging studies. Cerebral Cortex, 24(11), 2981–2990. https://doi.org/10.1093/cercor/bht154

Camras, L. A. (2011). Differentiation, dynamical integration and functional emotional development. Emotion Review, 3(2), 138–146. https://doi.org/10.1177/1754073910387944

Casey, B. J. (2015). Beyond simple models of self-control to circuit-based accounts of adolescent behavior. Annual Review of Psychology, 66, 295–319. https://doi.org/10.1146/annurev-psych-010814-015156

Chang, L. J., Gianaros, P. J., Manuck, S. B., & Krishnan, A. (2015). A sensitive and specific neural signature for picture-induced negative affect. PLOS Biology, 13(6), 1–28. https://doi.org/10.1371/journal.pbio.1002180

Choudhury, S., Blakemore, S. J., & Charman, T. (2006). Social cognitive development during adolescence. Social Cognitive and Affective Neuroscience, 1(3), 165–174. https://doi.org/10.1093/scan/nsl024

Christophel, T. B., Iamshchinina, P., Yan, C., Allefeld, C., & Haynes, J.-D. (2018). Cortical specialization for attended versus unattended working memory. Nature Neuroscience. https://doi.org/10.1038/s41593-018-0094-4

Cole, P. M., & Deater-Deckard, K. (2009). Emotion regulation, risk, and psychopathology. Journal of Child Psychology and Psychiatry and Allied Disciplines, 50(11), 1327–1330. https://doi.org/10.1111/j.1469-7610.2009.02180.x

Cole, P. M., Martin, S. E., & Dennis, T. A. (2004). Emotion regulation as a scientific construct: Methodological challenges and directions for child development research. Child Development, 75(2), 317–333.

Cole, S. R., & Voytek, B. (2017). Brain oscillations and the importance of waveform shape. Trends in Cognitive Sciences, 21(2), 137–149. https://doi.org/10.1016/j.tics.2016.12.008

Cole, S. R., & Voytek, B. (2018). Cycle-by-cycle analysis of neural oscillations. BioRxiv, 1–28.

Crone, E. A., & van der Molen, M. W. (2004). Developmental changes in real life decision making: Performance on a gambling task previously shown to depend on the ventromedial prefrontal cortex. Developmental Neuropsychology, 25(3), 251–279. https://doi.org/10.1207/s15326942dn2503

Dehaene-Lambertz, G., Monzalvo, K., & Dehaene, S. (2018). The emergence of the visual word form: Longitudinal evolution of category-specific ventral visual areas during reading acquisition. PLOS Biology, 1–34. https://doi.org/10.1371/journal.pbio.2004103

Denny, B. T., & Ochsner, K. N. (2014). Behavioral effects of longitudinal training in cognitive reappraisal. Emotion, 14(2), 425–433. https://doi.org/10.1037/a0035276.Behavioral

Dinstein, I., Heeger, D. J., & Behrmann, M. (2015). Neural variability: Friend or foe? Trends in Cognitive Sciences, 19(6), 322–328. https://doi.org/10.1016/j.tics.2015.04.005

DuPre, E., & Spreng, R. N. (2017). Structural covariance networks across the lifespan, from 6-94 years of age. Network Neuroscience, 1–38. https://doi.org/10.1162/NETN_a_00016

Durston, S., Davidson, M. C., Tottenham, N., Galvan, A., Spicer, J., Fossella, J. A., & Casey, B. J. (2006). A shift from diffuse to focal cortical activity with development. Developmental Science, 9(1), 1–8. https://doi.org/10.1111/j.1467-7687.2005.00454.x

Eatough, E. M., Shirtcliff, E. A., Hanson, J. L., & Pollak, S. D. (2009). Hormonal reactivity to MRI scanning in adolescents. Psychoneuroendocrinology, 34(8), 1242–1246. https://doi.org/10.1111/j.1743-6109.2008.01122.x.Endothelial

Eisenberg, I., Bissett, P., Enkavi, A. Z., Li, J., MacKinnon, D., Marsch, L., & Poldrack, R. (2018). Uncovering mental structure through data-driven ontology discovery. PsyArXiv, 1–10. https://doi.org/10.31234/OSF.IO/FVQEJ

Eisenberg, N., Spinrad, T. L., & Knafo-Noam, A. (1998). Prosocial Development. John Wiley & Sons. https://doi.org/10.1002/9781118963418.childpsy315

Etkin, A., Büchel, C., & Gross, J. J. (2015). The neural bases of emotion regulation. Nature Reviews Neuroscience, 16(11), 693–700. https://doi.org/10.1038/nrn4044

Etzel, J. A., Zacks, J. M., & Braver, T. S. (2013). Searchlight analysis: Promise, pitfalls, and potential. NeuroImage, 78, 261–269. https://doi.org/10.1016/j.actbio.2009.04.013.Role

Foulkes, L., & Blakemore, S. J. (2018). Studying individual differences in human adolescent brain development. Nature Neuroscience, 1–9. https://doi.org/10.1038/s41593-018-0078-4

Galván, A., Van Leijenhorst, L., & McGlennen, K. M. (2012). Considerations for imaging the adolescent brain. Developmental Cognitive Neuroscience, 2(3), 293–302. https://doi.org/10.1016/j.dcn.2012.02.002

Garrett, D. D., Kovacevic, N., Mcintosh, A. R., & Grady, C. L. (2010). Blood oxygen level-dependent signal variability is more than just noise. The Journal of Neuroscience, 30(14), 4914–4921. https://doi.org/10.1523/JNEUROSCI.5166-09.2010

Garrett, D. D., Kovacevic, N., McIntosh, A. R., & Grady, C. L. (2011). The importance of being variable. Journal of Neuroscience, 31(12), 4496–4503. https://doi.org/10.1523/JNEUROSCI.5641-10.2011

Giuliani, N. R., & Pfeifer, J. H. (2015). Age-related changes in reappraisal of appetitive cravings during adolescence. NeuroImage, 108, 173–181. https://doi.org/10.1016/j.neuroimage.2014.12.037

Goldin, P. R., McRae, K., Ramel, W., & Gross, J. J. (2008). The neural bases of emotion regulation: Reappraisal and suppression of negative emotion. Biological Psychiatry, 63(6), 577–586. https://doi.org/10.1016/j.biopsych.2007.05.031

Gross, J. J. (2015). Emotion regulation: Current status and future prospects. Psychological Inquiry, 26(1), 1–26. https://doi.org/10.1080/1047840X.2014.940781

Gross, J. J., & Feldman Barrett, L. (2011). Emotion generation and emotion regulation: One or two depends on your point of view. Emotion Review, 3(1), 8–16. https://doi.org/10.1177/1754073910380974

Guassi Moreira, J. F., & Silvers, J. A. (2018). In due time: Neurodevelopmental considerations in the study of emotion regulation. In P. M. Cole & T. Hollenstein (Eds.), Emotion Regulation: A Matter of Time (pp. 111–134). New York: Routledge.

Guest, O., & Love, B. C. (2017). What the success of brain imaging implies about the neural code. ELife, 6, 1–16. https://doi.org/10.7554/eLife.21397

Haines, S. J., Gleeson, J., Kuppens, P., Hollenstein, T., Ciarrochi, J., Labuschagne, I., … Koval, P. (2016). The wisdom to know the difference: Strategy-situation fit in emotion regulation in daily life is associated with well-being. Psychological Science, 27(12), 1651–1659. https://doi.org/10.1177/0956797616669086

Haxby, J. V., Gobbini, M. I., Furey, M. L., Ishai, A., Schouten, J. L., & Pietrini, P. (2001). Distrubuted and overlapping representations of face and objects in ventral temporal cortex. Science, 293(5539), 2425–2430. https://doi.org/10.1126/science.1063736

Heathers, J. A. J., & Goodwin, M. S. (2017). Dead science in live psychology: A case study from heart rate variability (HRV). PsyArXiv, 1–34.

Heller, A. S., & Casey, B. J. (2016). The neurodynamics of emotion: Delineating typical and atypical emotional processes during adolescence. Developmental Science, 19(1), 3–18. https://doi.org/10.1111/desc.12373

Huth, A. G., Heer, W. A. De, Griffiths, T. L., Theunissen, F. E., & Gallant, J. L. (2016). Natural speech reveals the semantic maps that tile human cerebral cortex. Nature, 532(7600), 453–458. https://doi.org/10.1038/nature17637.Natural

Huth, A. G., Nishimoto, S., Vu, A. T., & Gallant, J. L. (2012). A continuous semantic space describes the representation of thousands of object and action categories across the human brain. Neuron, 76(6), 1210–1224. https://doi.org/10.1016/j.neuron.2012.10.014

Kim, J. H., & Richardson, R. (2010). New findings on extinction of conditioned fear early in development: Theoretical and clinical implications. Biological Psychiatry, 67(4), 297–303. https://doi.org/10.1016/j.biopsych.2009.09.003

Koolschijn, P. C. M. P., Schel, M. A., de Rooij, M., Rombouts, S. A. R. B., & Crone, E. A. (2011). A three-year longitudinal functional magnetic resonance imaging study of performance monitoring and test-retest reliability from childhood to early adulthood. Journal of Neuroscience, 31(11), 4204–4212. https://doi.org/10.1523/JNEUROSCI.6415-10.2011

Kriegeskorte, N., Cusack, R., & Bandettini, P. (2010). How does an fMRI voxel sample the neuronal activity pattern: Compact-kernel or complex spatiotemporal filter? NeuroImage, 49(3), 1965–1976. https://doi.org/10.1016/j.neuroimage.2009.09.059

Kriegeskorte, N., Mur, M., Ruff, D. A., Kiani, R., Bodurka, J., Esteky, H., … Bandettini, P. A. (2008). Matching categorical object representations in inferior temporal cortex of man and monkey. Neuron, 60(6), 1126–1141. https://doi.org/10.1016/j.neuron.2008.10.043

Kriegeskorte, N., Simmons, W. K., Bellgowan, P. S., & Baker, C. I. (2009). Circular analysis in systems neuroscience: The dangers of double dipping. Nature Neuroscience, 12(5), 535–540. https://doi.org/10.1038/nn.2303

Kurby, C. A., & Zacks, J. M. (2008). Segmentation in the perception and memory of events. Trends in Cognitive Sciences, 12(2), 72–79. https://doi.org/10.1016/j.tics.2007.11.004

Lang, P. J., Bradley, M. M., & Cuthbert, B. N. (2008). International affective picture system (IAPS): Affective ratings of pictures and instruction manual. Technical Report A-8. https://doi.org/10.1016/j.epsr.2006.03.016

Larsen, R. J., & Prizmic-Larsen, Z. (2006). Measuring emotions: Implications of a multimethod perspective. In M. Eid & E. Diener (Eds.), Handbook of multimethod measurement in psychology (pp. 337–351). Washington, D.C.: American Psychological Association.

Larson, R., Csikszentmihalyi, M., & Graef, R. (1980). Mood variability and the psycho-social adjustment of adolescents. Journal of Youth & Adolescence, 9(6), 469–490. https://doi.org/10.1007/978-94-017-9094-9_15

Leech, R., Scott, G., Carhart-Harris, R., Turkheimer, F., Taylor-Robinson, S. D., & Sharp, D. J. (2014). Spatial dependencies between large-scale brain networks. PLoS ONE, 9(6). https://doi.org/10.1371/journal.pone.0098500

Levesque, J., Joanette, Y., Mensour, B., Beaudoin, G., Leroux, J. M., Bourgouin, P., & Beauregard, M. (2004). Neural basis of emotional self-regulation in childhood. Neuroscience, 129(2), 361–369.

Luciana, M., Wahlstrom, D., Porter, J. N., & Collins, P. F. (2012). Dopaminergic modulation of incentive motivation in adolescence: Age-related changes in signaling, individual differences, and implications for the development of self-regulation. Developmental Psychology, 48(3), 844–861. https://doi.org/10.1037/a0027432.Dopaminergic

McLaughlin, K. A., Garrard, M. C., & Somerville, L. H. (2015). What develops during emotional development? A component process approach to identifying sources of psychopathology risk in adolescence. Dialogues in Clinical Neuroscience, 17(4), 403–410.

McLaughlin, K. A., Peverill, M., Gold, A. L., Alves, S., & Sheridan, M. A. (2015). Child maltreatment and neural systems underlying emotion regulation. Journal of the American Academy of Child and Adolescent Psychiatry, 54(9), 753–62. https://doi.org/10.1016/j.jaac.2015.06.010

McRae, K., Gross, J. J., Weber, J., Robertson, E. R., Sokol-Hessner, P., Ray, R. D., … Ochsner, K. N. (2012). The development of emotion regulation: An fMRI study of cognitive reappraisal in children, adolescents and young adults. Social Cognitive and Affective Neuroscience, 7(1), 11–22. https://doi.org/10.1093/scan/nsr093

Mills, K. L., Goddings, A. L., Clasen, L. S., Giedd, J. N., & Blakemore, S. J. (2014). The developmental mismatch in structural brain maturation during adolescence. Developmental Neuroscience, 36(3–4), 147–160. https://doi.org/10.1159/000362328

Mills, K. L., Lalonde, F., Clasen, L. S., Giedd, J. N., & Blakemore, S. J. (2014). Developmental changes in the structure of the social brain in late childhood and adolescence. Social Cognitive and Affective Neuroscience, 9(1), 123–131. https://doi.org/10.1093/scan/nss113

Mischel, H. N., & Mischel, W. (1983). The development of children’s knowledge of self-control strategies. Child Development, 54(3), 603. https://doi.org/10.2307/1130047

Mischel, W., & Baker, N. (1975). Cognitive appraisals and transformations in delay behavior. Journal of Personality and Social Psychology, 31(2), 254–261. https://doi.org/10.1037/h0076272

Mumford, J. A., Davis, T., & Poldrack, R. A. (2014). The impact of study design on pattern estimation for single-trial multivariate pattern analysis. NeuroImage, 103, 130–138. https://doi.org/10.1016/j.neuroimage.2014.09.026

Mumford, J. A., Turner, B. O., Ashby, F. G., & Poldrack, R. A. (2012). Deconvolving BOLD activation in event-related designs for multivoxel pattern classification analyses. NeuroImage, 59(3), 2636–2643. https://doi.org/10.1016/j.neuroimage.2011.08.076

Nomi, J. S., Bolt, T. S., Ezie, C. E. C., Uddin, L. Q., & Heller, A. S. (2017). Moment-to-moment BOLD Signal variability reflects regional changes in neural flexibility across the lifespan. The Journal of Neuroscience, 37(22), 5539–5548. https://doi.org/10.1523/JNEUROSCI.3408-16.2017

Northoff, G., & Bermpohl, F. (2004). Cortical midline structures and the self. Trends in Cognitive Sciences, 8(3), 102–107. https://doi.org/10.1016/j.tics.2004.01.004

Ochsner, K. N., & Gross, J. J. (2005). The cognitive control of emotion. Trends in Cognitive Sciences.

Ochsner, K. N., & Gross, J. J. (2008). Insights from social cognitive and affective neuroscience. Current Directions in Psychological Science, 17(2), 153–158. https://doi.org/10.1111/j.1467-8721.2008.00566.x

Ochsner, K. N., Ray, R. D., Cooper, J. C., Robertson, E. R., Chopra, S., Gabrieli, J. D. E., & Gross, J. J. (2004). For better or for worse: Neural systems supporting the cognitive down- and up-regulation of negative emotion. NeuroImage, 23(2), 483–499. https://doi.org/10.1016/j.neuroimage.2004.06.030

Ochsner, K. N., Silvers, J. A., & Buhle, J. T. (2012). Review and evolving model of the cognitive control of emotion. Annals of the New York Academy of Sciences, (1251), E1–E24. https://doi.org/10.1111/j.1749-6632.2012.06751.x.Functional

Oosterhof, N. N., Connolly, A. C., & Haxby, J. V. (2016). CoSMoMVPA: Multi-modal multivariate pattern analysis of neuroimaging data in Matlab/GNU Octave. Frontiers in Neuroinformatics, 10, 1–27. https://doi.org/10.3389/fninf.2016.00027

Ordaz, S. J., Foran, W., Velanova, K., & Luna, B. (2013). Longitudinal growth curves of brain function underlying inhibitory control through adolescence. The Journal of Neuroscience, 33(46), 18109–18124. https://doi.org/10.1523/JNEUROSCI.1741-13.2013

Panno, A., Lauriola, M., & Figner, B. (2013). Emotion regulation and risk taking: Predicting risky choice in deliberative decision making. Cognition & Emotion, 27(2), 326–34. https://doi.org/10.1080/02699931.2012.707642

Park, D. C., Polk, T. A., Park, R., Minear, M., Savage, A., & Smith, M. R. (2004). Aging reduces neural specialization in ventral visual cortex. Proceedings of the National Academy of Sciences, 101(35), 13091–13095. https://doi.org/10.1073/pnas.0405148101

Patel, G. H., Kaplan, D. M., & Snyder, L. H. (2014). Topographic organization in the brain: Searching for general principles. Trends in Cognitive Sciences, 18(7), 351–363. https://doi.org/10.1016/j.tics.2014.03.008

Petroni, A., Cohen, S. S., Ai, L., Langer, N., Henin, S., Vanderwal, T., … Parra, L. C. (2018). The variability of neural responses to naturalistic videos change with age and sex. Eneuro, 5(February), ENEURO.0244-17.2017. https://doi.org/10.1523/ENEURO.0244-17.2017

Pfeifer, J. H., & Blakemore, S. J. (2012). Adolescent social cognitive and affective neuroscience: Past, present, and future. Social Cognitive and Affective Neuroscience, 7(1), 1–10. https://doi.org/10.1093/scan/nsr099

Pfeifer, J. H., & Peake, S. J. (2012). Self-development: Integrating cognitive, socioemotional, and neuroimaging perspectives. Developmental Cognitive Neuroscience, 2(1), 55–69. https://doi.org/10.1016/j.dcn.2011.07.012

Phan, K. L., Fitzgerald, D. A., Nathan, P. J., Moore, G. J., Uhde, T. W., & Tancer, M. E. (2005). Neural substrates for voluntary suppression of negative affect: A functional magnetic resonance imaging study. Biological Psychiatry, 57(3), 210–219. https://doi.org/10.1016/j.biopsych.2004.10.030

Pinneo, L. R. (1966). On noise in the nervous system. Psychological Review, 73(3), 242–247.

Poldrack, R. A. (2015). Is “efficiency” a useful concept in cognitive neuroscience? Developmental Cognitive Neuroscience, 11, 12–17. https://doi.org/10.1016/j.dcn.2014.06.001

Polk, T. A., Stallcup, M., Aguirre, G. K., Alsop, D. C., D’Esposito, M., Detre, J. A., & Farah, M. J. (2002). Neural specialization for letter recognition. Journal of Cognitive Neuroscience, 14(2), 145–159. https://doi.org/10.1162/089892902317236803

Pyatt, G. (1976). On the interpretation and disaggregation of gini coefficients. The Economic Journal, 86(342), 243–255.

Ray, R. D., McRae, K., Ochsner, K. N., & Gross, J. J. (2010). Cognitive reappraisal of negative affect: Converging evidence from EMG and self-report. Emotion, 10(4), 587–592. https://doi.org/10.1016/j.pmrj.2014.02.014.Lumbar

Richardson, H., Lisandrelli, G., Riobueno-Naylor, A., & Saxe, R. (2018). Development of the social brain from age three to twelve years. Nature Communications, 9(1), 1027. https://doi.org/10.1038/s41467-018-03399-2

Richmond, L. L., & Zacks, J. M. (2017). Constructing Experience: Event Models from Perception to Action. Trends in Cognitive Sciences, 21(12), 962–980. https://doi.org/10.1016/j.tics.2017.08.005

Rissman, J., Gazzaley, A., & D’Esposito, M. (2004). Measuring functional connectivity during distinct stages of a cognitive task. NeuroImage, 23(2), 752–763. https://doi.org/10.1016/j.neuroimage.2004.06.035

Roalf, D. R., Gur, R. E., Ruparel, K., Calkins, M., Satterthwaite, T., Bilker, W. B., … Ruben, C. (2014). Within-individual variability in neurocognitive performance: age and sex-related differences in children and youths from ages 8 to 21. Neuropsychology, 28(4), 506–518. https://doi.org/10.1037/neu0000067.Within-Individual

Seghier, M. L., & Price, C. J. (2018). Interpreting and utilising intersubject variability in brain function. Trends in Cognitive Sciences, 22(6), 517–530. https://doi.org/10.1016/j.tics.2018.03.003

Siegel, J. S., Power, J. D., Dubis, J. W., Vogel, A. C., Church, J. A., Schlaggar, B. L., & Petersen, S. E. (2014). Statistical improvements in functional magnetic resonance imaging analyses produced by censoring high-motion data points. Human Brain Mapping, 35(5), 1981–1996. https://doi.org/10.1002/hbm.22307.Statistical

Siegler, R. S. (1994). Cognitive Variability : A Key to Understanding Cognitive Development. Current Directions in Psychological Science, 3(1), 1–5.

Siegler, R. S. (2007). Cognitive variability. Developmental Science, 10(1), 104–109. https://doi.org/10.1111/j.1467-7687.2007.00571.x

Silvers, J. A., Insel, C., Powers, A., Franz, P., Helion, C., Martin, R. E., … Ochsner, K. N. (2016). vlPFC – vmPFC – amygdala interactions underlie age-related fifferences in cognitive regulation of emotion. Cerebral Cortex, ([Epub Ahead of Print]), 1–13. https://doi.org/10.1093/cercor/bhw073

Silvers, J. A., Insel, C., Powers, A., Franz, P., Weber, J., Mischel, W., … Ochsner, K. N. (2014). Curbing craving: Behavioral and brain evidence that children regulate craving when instructed to do so but have higher baseline craving than adults. Psychological Science, 25(10), 1932–1942. https://doi.org/10.1177/0956797614546001

Silvers, J. A., McRae, K., Gabrieli, J. D. E., Gross, J. J., Remy, K. A., & Ochsner, K. N. (2012). Age-related differences in emotional reactivity, regulation, and rejection sensitivity in adolescence. Emotion, 12(6), 1235–1247. https://doi.org/10.1037/a0028297.Age-Related

Silvers, J. A., Shu, J., Hubbard, A. D., Weber, J., & Ochsner, K. N. (2015). Concurrent and lasting effects of emotion regulation on amygdala response in adolescence and young adulthood. Developmental Science, 18(5), 771–784. https://doi.org/10.1111/desc.12260

Smith, L. B., & Thelen, E. (2003). Development as a dynamic system. Trends in Cognitive Sciences, 7(8), 343–348. https://doi.org/10.1016/S1364-6613(03)00156-6

Somerville, L. H., Jones, R. M., & Casey, B. J. (2010). A time of change: Behavioral and neural correlates of adolescent sensitivity to appetitive and aversive environmental cues. Brain and Cognition, 72(1), 124–133. https://doi.org/10.1016/j.bandc.2009.07.003

Somerville, L. H., Jones, R. M., Ruberry, E. J., Dyke, J. P., Glover, G., & Casey, B. J. (2013). The medial prefrontal cortex and the emergence of self-conscious emotion in adolescence. Psychological Science, 24(8), 1554–1562. https://doi.org/10.1177/0956797613475633

Sowell, E. R., Thompson, P. M., Tessner, K. D., & Toga, A. W. (2001). Mapping continued brain growth and gray matter density reduction in dorsal frontal cortex: Inverse relationships during postadolescent brain maturation. The Journal of Neuroscience, 21(22), 8819–29. https://doi.org/21/22/8819 [pii]

Thompson, R., & Goodman, M. (2010). Development of emotion regulation. In A. M. Kring & D. M. Sloan (Eds.), Emotion Regulation and Psychopathology: A transdiagnostic Approach to Etiology and Treatment (pp. 38–58). New York, NY: The Guilford Press.

Vij, S. G., Nomi, J. S., Dajani, D. R., & Uddin, L. Q. (2018). Evolution of spatial and temporal features of functional brain networks across the lifespan. NeuroImage. https://doi.org/10.1016/j.neuroimage.2018.02.066

